# Ceramide-1-phosphate transfer protein enhances lipid transport by disrupting hydrophobic lipid–membrane contacts

**DOI:** 10.1101/2022.09.10.507427

**Authors:** Julia R. Rogers, Phillip L. Geissler

## Abstract

Cellular distributions of the sphingolipid ceramide-1-phosphate (C1P) impact essential biological processes. C1P levels are spatiotemporally regulated by ceramide-1-phosphate transfer protein (CPTP), which efficiently shuttles C1P between organelle membranes. Yet, how CPTP rapidly extracts and inserts C1P into a membrane remains unknown. Here, we devise a multiscale simulation approach to elucidate biophysical details of CPTP-mediated C1P transport. We find that CPTP binds a membrane poised to extract and insert C1P and that membrane binding promotes conformational changes in CPTP that facilitate C1P uptake and release. By significantly disrupting a lipid’s local hydrophobic environment in the membrane, CPTP lowers the activation free energy barrier for passive C1P desorption and enhances C1P extraction from the membrane. Upon uptake of C1P, further conformational changes may aid membrane unbinding in a manner reminiscent of the electrostatic switching mechanism used by other lipid transfer proteins. Insertion of C1P into an acceptor membrane, eased by a decrease in membrane order by CPTP, restarts the transfer cycle. Most notably, we provide molecular evidence for CPTP’s ability to catalyze C1P extraction by breaking hydrophobic C1P–membrane contacts with compensatory hydrophobic lipid–protein contacts. Our work, thus, provides biophysical insights into how CPTP efficiently traffics C1P between membranes to maintain sphingolipid homeostasis and, additionally, presents a simulation method aptly suited for uncovering the catalytic mechanisms of other lipid transfer proteins.

**Author summary:** Critical cellular processes require spatiotemporal regulation of sphingolipid levels among organelle membranes. Programmed cell death and inflammation, for example, are impacted by the distribution of ceramide-1-phosphate (C1P). C1P levels are specifically altered by ceramide-1-phosphate transfer protein (CPTP), which mediates C1P intermembrane transport. Using a multiscale simulation approach tailored to studying lipid transport, we elucidate key steps in the molecular mechanism used by CPTP to rapidly transport C1P between membranes: Through conformational changes that are coupled to membrane binding, CPTP significantly disrupts C1P’s local hydrophobic environment in a membrane and catalyzes its extraction. Since this catalytic mechanism is biophysically related to that of passive lipid transport, it may be ubiquitously used by lipid transport proteins to rapidly traffic lipids between membranes and ensure membrane homeostasis. Our multiscale simulation approach offers a framework to test this hypothesis and, thus, further our molecular knowledge of how lipid transfer proteins function to regulate cellular lipid distributions.

## Introduction

Sphingolipids play critical roles in numerous cellular processes, including cell growth, differentiation, migration, and survival [1, 2]. Normal biological function requires spatiotemporal regulation of sphingolipid levels among organelle membranes [2– 4], and altered distributions are linked to the pathophysiology of diseases including cancers, inflammatory diseases, metabolic syndromes, and neurological disorders [1, 5–7]. Precise sphingolipid distributions are maintained through coordinated regulation of sphingolipid metabolism within specific organelle membranes and transport between them [3, 4, 8, 9]. While interorganelle transport can occur through vesicular trafficking [3, 9], individual sphingolipids are also specifically transported between membranes by lipid transfer proteins [9–14]. One such protein, ceramide-1-phosphate transfer protein (CPTP) transports ceramide-1-phosphate (C1P), a mitogenic and prosurvival sphingolipid [1, 15], from the *trans*-Golgi, where it’s synthesized from ceramide, to nuclear and plasma membranes [16]. By selectively trafficking C1P from the *trans*-Golgi, CPTP modulates pro-inflammatory eicosanoid production, interluekin release, and autophagy, and, thus, may help reduce the risk of chronic inflammation [16–18].

Efforts to fully decipher and modulate CPTP’s role in both homeostasis and disease, nevertheless, remain hampered by an incomplete biophysical understanding of how CPTP rapidly traffics C1P between organelles. Structural and biochemical studies [16, 19, 20] have elucidated initial details about how CPTP functions as a cytosolic lipid shuttle [11, 14]: During transit, CPTP houses C1P within a hydrophobic cavity at the interior of its all α-helical, sandwich-like structure [16], thereby shielding C1P’s hydrophobic tails from unfavorable exposure to solvent. While this partially explains why CPTP facilitated transport of C1P is more rapid than passive transport, it fails to explain how CPTP swiftly extracts (and inserts) C1P from (into) a membrane. Based on crystallographic measurements, a cleft-like gating mechanism has been proposed to control access to CPTP’s hydrophobic cavity and to facilitate C1P entry and exit [16]; however, because such molecular motions have yet to be resolved, their importance for C1P uptake and release is unclear. An additional puzzle is how CPTP overcomes the significant free energy barrier for lipid desorption from a membrane, which limits passive lipid transport rates [21–25]. We recently demonstrated that the activation free energy barrier for passive lipid desorption reflects the thermodynamic cost of breaking hydrophobic lipid–membrane contacts [25], and that it can be lowered by decreasing the hydrophobicity of the transferring lipid’s local membrane environment [26]. Thus, we hypothesize that CPTP rapidly extracts C1P from a membrane by disrupting hydrophobic C1P–membrane contacts, thereby lowering the free energy barrier for lipid desorption and catalyzing C1P transport.

Here, we investigate how CPTP performs individual steps of the C1P transfer cycle using a multiscale molecular dynamics (MD) simulation approach. Significant structural differences between the apo and C1P-bound forms, observed in both published crystal structures [16] and our solution-phase simulations, indicate that conformational changes are required to accommodate C1P in CPTP’s hydrophobic cavity. Consistent with previous results [16, 27], CPTP exhibits a single favorable membrane binding pose that aptly positions the entrance to its hydrophobic cavity at the membrane interface in our simulations. We find that membrane binding promotes rearrangements of CPTP’s helices as expected for a cleft-like gating mechanism to facilitate C1P entry and exit into its hydrophobic cavity. Key to CPTP’s catalytic function, we find that the apo form significantly disrupts a lipid’s local hydrophobic environment in the membrane. As a result, CPTP reduces the rate-limiting free energy barrier for C1P desorption. We suggest that uptake of C1P may then trigger conformational changes that prompt membrane unbinding and continuation of the transfer cycle. Overall, our simulations shed light on how CPTP rapidly traffics C1P between membranes and also offer molecular explanations for previous experimental findings.

## Results

We devise a multiscale simulation approach that builds upon previous structural studies [16, 27] and our earlier studies of passive lipid transport [25, 26] to efficiently and accurately characterize key elementary steps of C1P transport by CPTP. Specifically, we model the transport of C1P with a saturated, 16-carbon acyl chain, which is CPTP’s most likely *in vivo* substrate [16], between membranes composed of 1-palmitoyl-2-oleoyl-*sn*-glycero-3-PC (POPC), which have been used in experimental transfer assays [16, 19, 20], by human CPTP. We use atomistic simulations to resolve conformational changes that occur upon membrane binding and to capture C1P extraction from and insertion into a membrane, whereas we use coarse-grained simulations to observe multiple membrane binding events, which require microsecond-long simulations. Using an enhanced sampling method specifically targeting the degrees of freedom that hinder passive lipid transport, we calculate an all-atom free energy profile of CPTP-mediated C1P extraction and compare it to that of passive C1P desorption from a membrane to assess CPTP’s catalytic abilities.

### CPTP’s hydrophobic cavity must expand to accommodate C1P

CPTP has a two-layered, all *α*-helical topology that is homologous with the glycolipid transfer protein (GLTP) fold [18, 28–30]. Within the interior of CPTP’s sandwich-like structure composed of nine *α*-helices is a hydrophobic cavity that adaptively expands to accommodate C1P. We first examined how the size of the cavity and overall structures of the apo and C1P-bound forms of CPTP differ in solution. To initialize solution-phase simulations, we used the most complete crystal structure of CPTP with C1P fully bound in its hydrophobic cavity (PDB 4K85). Final structures from our solution-phase simulations of the apo and C1P-bound forms are shown in Fig 1A-C. Without C1P present in the cavity, CPTP’s helices reorient themselves, closing off the entrance to the hydrophobic cavity and barring entry of solvent molecules. In particular, helix *α*2, which is located on one side of CPTP’s sandwich topology, and helices *α4* and *α*8, which are located on the opposing side, angle into the hydrophobic cavity and fill the space otherwise occupied by C1P (Fig 1A). Due to the closer packing of multiple helices in the apo form, the hydrophobic cavity collapses (Fig 1B and 1C). To quantify the change in hydrophobic cavity volume, we use the geometry-based detection algorithm MDpocket [31] to identify the void between CPTP’s two *α*-helical layers unoccupied by protein, and find that the total volume of the hydrophobic cavity in the apo form is on average only half the volume of the cavity in the C1P-bound form (Fig 1D). Consistent with conclusions made based on crystal structures [16], our results indicate that CPTP must undergo significant conformational changes to accommodate C1P in its hydrophobic cavity. Differences in the structures of the C1P-bound and apo forms in our simulations are further suggestive of the previously proposed cleft-like gating mechanism [16]: To accomodate C1P, helices on one side of CPTP’s sandwich-like structure are collectively displaced due to a substantial change in the location of helix *α*3, which connects residues on either side of the hydrophobic cavity (S1 Fig and S1 Text, Note 1).

**Fig 1.**
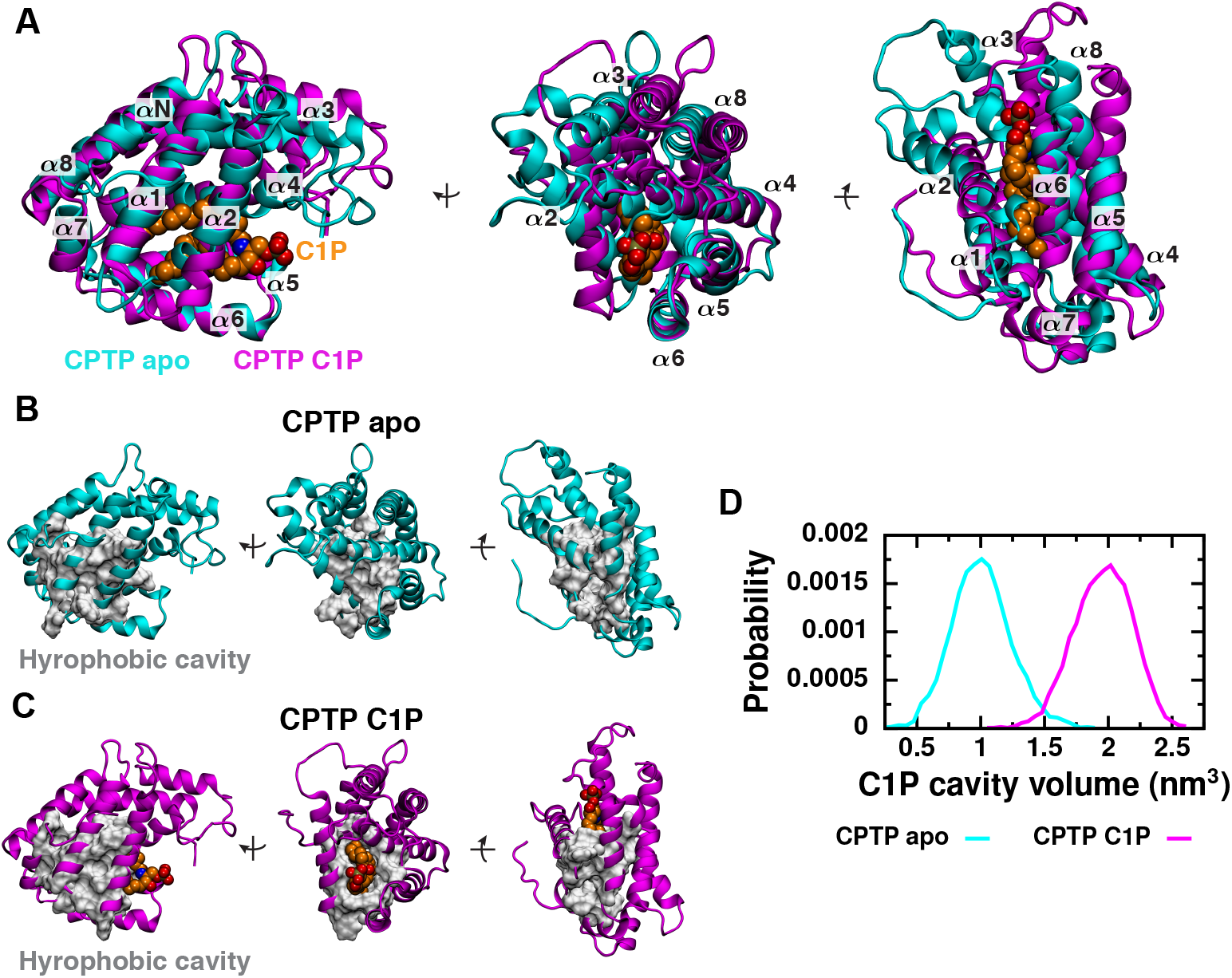
In solution, conformational differences between the apo and C1P-bound forms of CPTP tune the accessibility and volume of the hydrophobic cavity. (A) Superimposed structures of the apo form (cyan) and C1P-bound form (magenta) obtained from solution-phase simulations are shown from three views. Structures were aligned to minimize the root-mean-square deviation (RMSD) of C*α* atoms of helix *α*6. Helices (*α*N and *α*1 − *α*8) are numbered from amino (N) to carboxy (C) termini. C1P is rendered as van der Waals spheres and colored orange. (B and C) Residues lining the hydrophobic cavity are shown as a gray surface on the (B) apo and (C) C1P-bound structures. (D) Distributions of the total volume of the hydrophobic cavity during solution-phase simulations of the apo and C1P-bound forms. Cavity volume is calculated as the volume of CPTP’s interior unoccupied by protein heavy atoms using MDpocket [31] and includes volume occupied by C1P in the C1P-bound form.

### Membrane binding poises CPTP to extract C1P

We next characterized how CPTP, in both apo and C1P-bound forms, binds to a POPC membrane. To observe multiple spontaneous binding events, we performed coarse-grained simulations (S2 Fig). While the coarse-grained model accurately recapitulates the structure and internal dynamics of CPTP observed in our atomistic solution-phase simulations (S3 Fig and S4 Fig), it cannot capture conformational changes that may occur upon membrane binding. Thus, we converted membrane-bound, coarse-grained configurations back into all-atom representations in order to resolve conformational changes that may result from CPTP–membrane interactions.

A single binding pose is observed in coarse-grained simulations (Fig 2 and S2 Fig), and CPTP remains favorably bound to the membrane in the same overall pose in all-atom simulations (Fig 2). This membrane-bound pose matches that predicted based on CPTP’s crystal structure modeled in the presence of an implicit membrane [16], which has previously been shown to be stable in all-atom MD simulations [27]. In both apo and C1P-bound forms, the loop connecting helices *α*1 and *α*2 along with helix *α*6 inserts into the membrane’s hydrophobic core (Fig 2A). A combination of hydrophobic and aromatic residues, specifically I49, F50, and F52 at the N-terminus of helix *α*2 and W152, V153, and V160 of helix *α*6, anchor CPTP to the membrane as predicted from crystal structures [16] and identified in previous simulations [27]. When bound to the membrane, the entrance to CPTP’s hydrophobic cavity is aptly positioned at the membrane–solvent interface (Fig 2B and 2C). Additionally, the cleft between CPTP’s two *α*-helical layers faces into the membrane’s hydrophobic core, providing an avenue for C1P to move from the membrane into CPTP’s hydrophobic cavity without exposing its tails to solvent.

**Fig 2.**
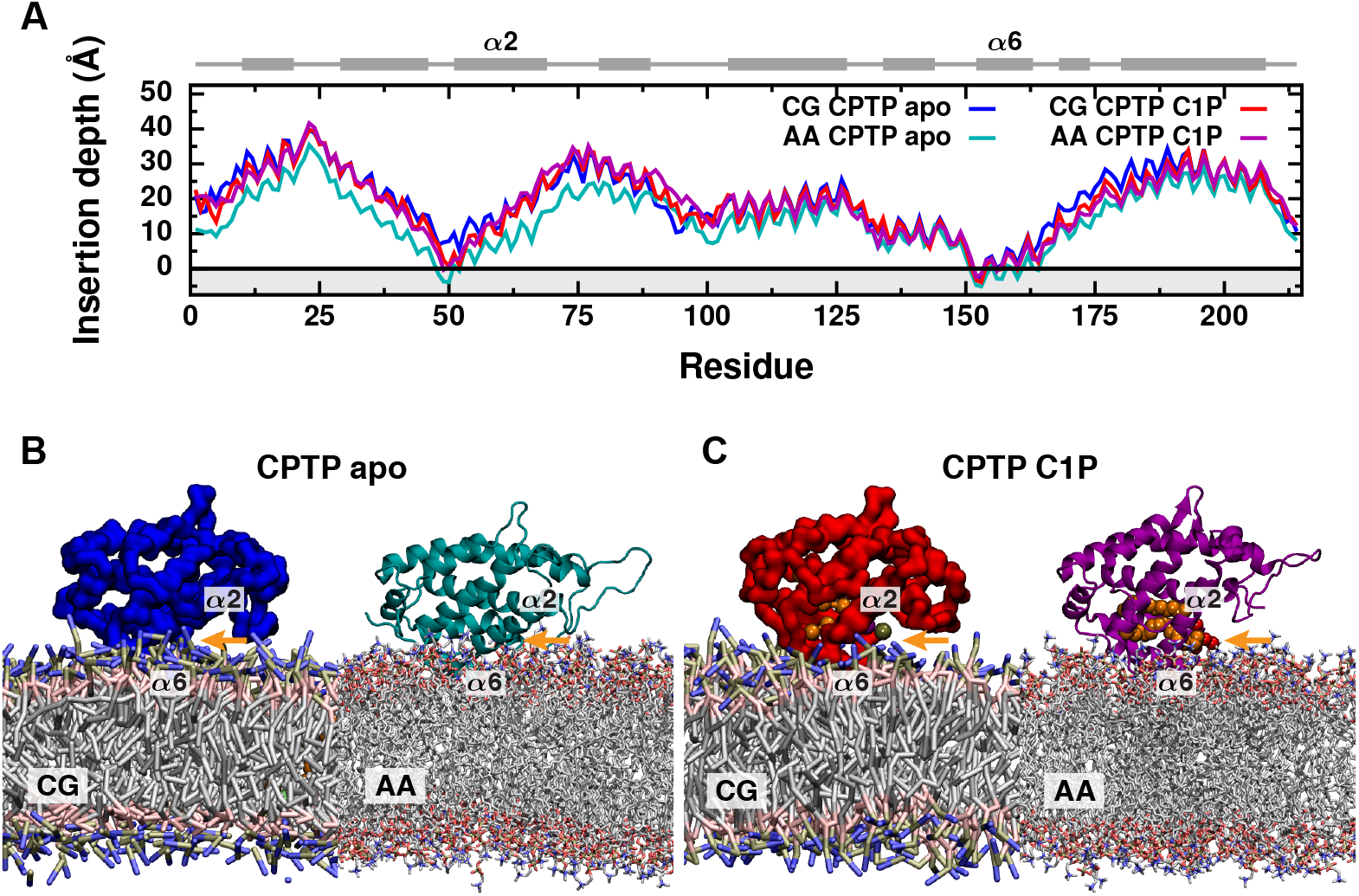
Both apo and C1P-bound forms of CPTP bind a membrane poised to extract or insert C1P. (A) Average insertion depth of each residue measured relative to the average position of lipid phosphate groups (black line) from coarse-grained (CG) and all-atom (AA) simulations. The gray region highlights the location of the membrane’s hydrophobic core. CPTP’s secondary structure is schematically illustrated above with helices represented as rectangles and unstructured loop regions as lines. (B and C) Representative configurations of the (B) apo form and (C) C1P-bound form of CPTP bound to a POPC membrane for both coarse-grained and all-atom representations. The entrance to CPTP’s hydrophobic cavity is indicated with an orange arrow. C1P is rendered as van der Waals spheres and colored orange.

Subtle differences in the binding pose of the apo form observed in all-atom versus coarse-grained representations (Fig 2A and 2B) suggest that conformational changes, which cannot be observed with the coarse-grained model, may be coupled to membrane binding. Indeed, closer inspection of the all-atom simulations indicates that interactions with the membrane promote initial widening of the entrance to CPTP’s hydrophobic cavity. Helix *α*2, which partially gates the entrance to the hydrophobic cavity (Fig 1A), is rotated outward in membrane-bound structures of both the apo and C1P-bound forms compared to their respective solution-phase structures (Fig 3). We quantify this conformational change by the polar angle *θ*_*α*2_ of helix *α*2 in a coordinate system defined by the membrane surface (Fig 3B). In order to calculate *θ*_*α*2_ for both solution-phase and membrane-bound structures, we use the plane that contains the top surface of helix *α*6 to approximate the plane of the membrane surface. We use helix *α*6 as an internal reference since helix *α*6 is approximately parallel to the membrane surface when CPTP is membrane bound (Fig 2A) and has a rather rigid structure (S3 Fig and S4 Fig). A decrease in *θ*_*α*2_ indicates opening of the gate created by helix *α*2. When C1P is bound, CPTP’s cavity is expanded and the gate widened (highlighted by orange arrows in Fig 3A), resulting in decreased *θ*_*α*2_ compared to the apo form (Fig 3C). As shown in Fig 3C, membrane-bound structures of CPTP have reduced *θ*_*α*2_ on average, corresponding to more open gates compared to their respective solution-phase structures. We draw similar conclusions from principal component analysis of CPTP’s structural ensemble (S1 Text, Note 1): Membrane-bound structures have more open helix *α*2 gates and wider clefts into CPTP’s hydrophobic cavity. By locking down the gate created by helix *α*2 when in solution, the hydrophobic cavity, and any lipid inside, remains sealed off from solvent. Initial widening of the entrance to the hydrophobic cavity upon membrane binding (highlighted by gray arrows in Fig 3A) helps ready CPTP to extract or insert C1P.

**Fig 3.**
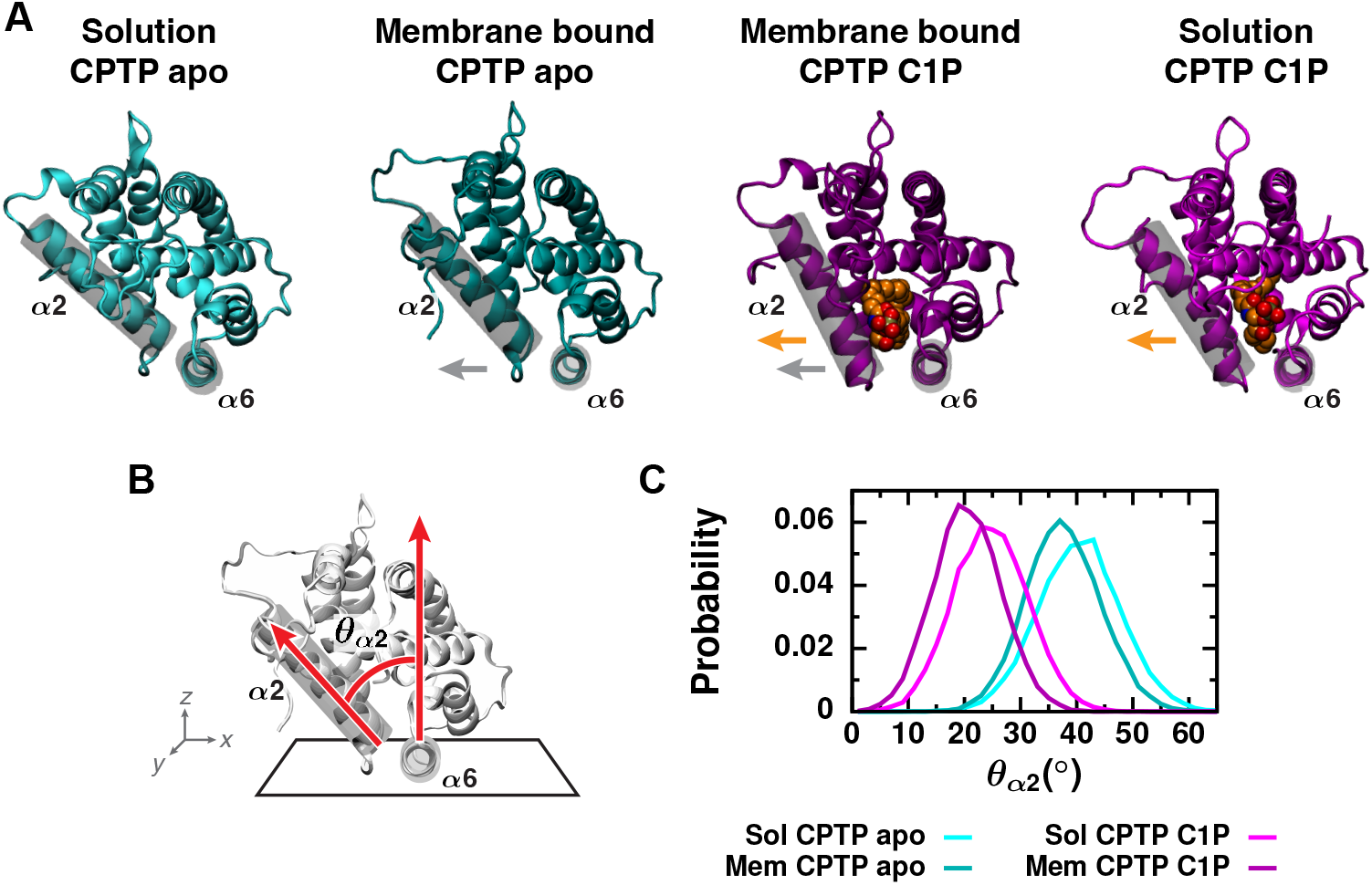
Conformational changes that promote the opening of gating helix α2 are coupled to membrane binding. (A) Orientation of helix *α*2 relative to helix *α*6 is highlighted on representative structures of both the apo and C1P-bound forms in both solution and bound to the membrane. Both helices are outlined by a gray cylinder. C1P is rendered as van der Waals spheres and colored orange. Orange arrows indicate gate opening upon C1P-binding to CPTP, and gray arrows indicate gate opening upon membrane binding. (B) Definition of angle used to characterize gate opening. A coordinate system is defined by the plane containing helix *α*6, which nearly corresponds to the location of the membrane–solvent interface when CPTP is membrane bound. *θ*_*α*2_ is the polar angle of helix *α*2. (C) Distributions of *θ*_*α*2_ from all-atom simulations of the apo and C1P-bound forms of CPTP both in solution and bound to the membrane.

Both the apo and C1P-bound forms of CPTP bind the membrane in the same overall pose and exhibit conformational changes of similar magnitude upon binding; however, such conformational changes enable the apo form to insert more deeply into the membrane’s hydrophobic core (Fig 2). In the apo form, a slight widening of the entrance to the hydrophobic cavity brings helix *α*2 closer to the membrane surface. As a result, energetically favorable interactions between residues on helix *α*2 and the membrane are present in the apo form but not the C1P-bound form (Fig 4). In particular, two positively charged residues, R62 and R66, form significantly favorable electrostatic interactions with the membrane and anchor CPTP in its apo but not C1P-bound form. By anchoring helix *α*2 to the membrane, such favorable interactions may modulate opening of the entrance to CPTP’s hydrophobic cavity in the apo form. Complete opening of the gate created by helix *α*2 to accommodate C1P inside CPTP’s hydrophobic cavity breaks these favorable electrostatic interactions with the membrane.

**Fig 4.**
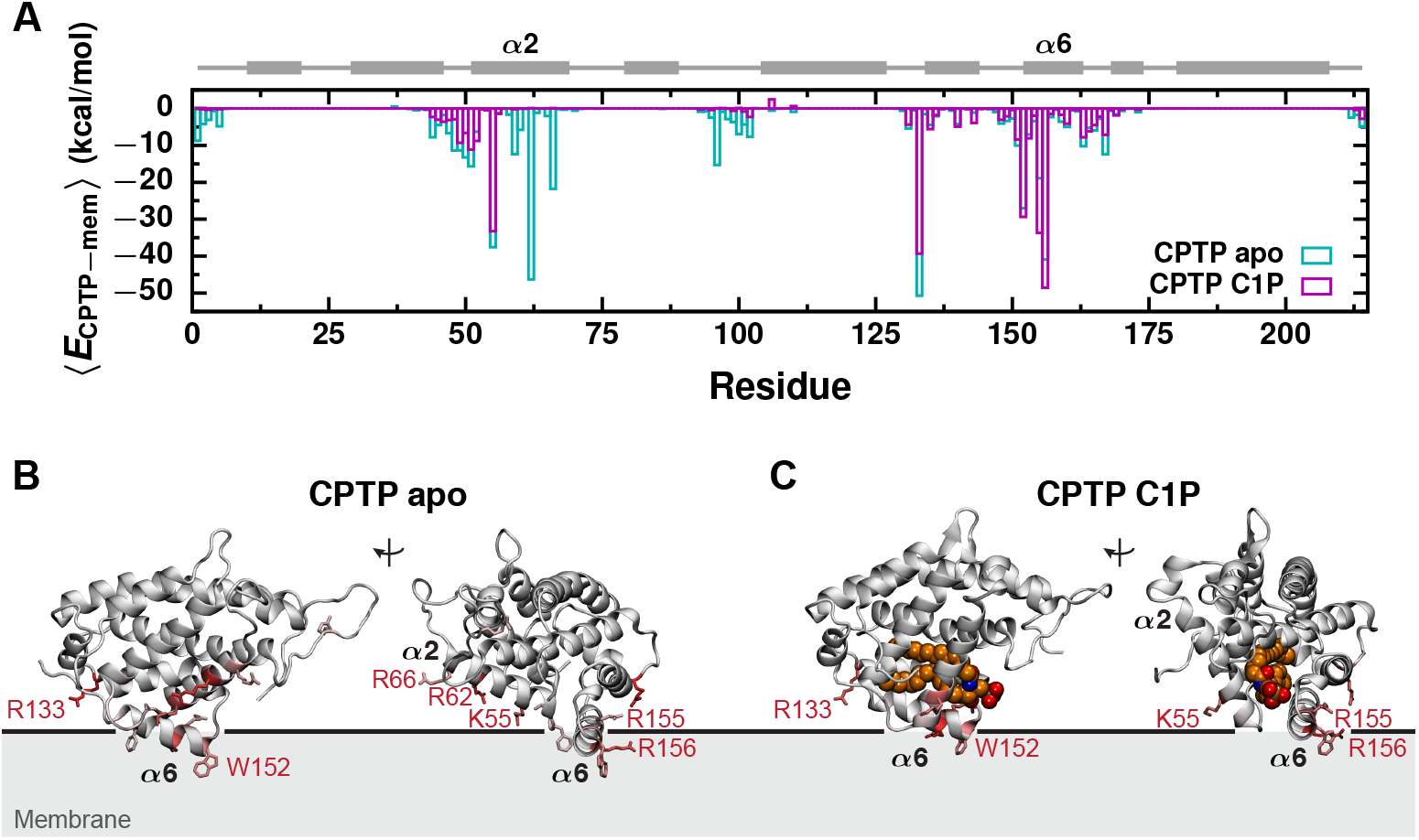
Favorable electrostatic interactions anchor the apo form deeply into the membrane. (A) Average interaction energy between each protein residue and the membrane, 〈*E*_CPTP–mem_〉, from all-atom simulations of the apo and C1P-bound forms of CPTP bound to a POPC membrane. CPTP’s secondary structure is schematically illustrated above with helices represented as rectangles and unstructured loop regions as lines. (B and C) 〈*E*_CPTP–mem_〉 mapped onto the structures of the (B) apo and (C) C1P-bound forms. 〈*E*_CPTP–mem_〉 is indicated by the color of the residue. Residues that most favorably interact with the membrane are colored dark red, those that weakly or do not interact with the membrane are colored gray. Side chains of residues with 〈*E*_CPTP–mem_〉 < −12 kcal/mol are rendered. Residues with 〈*E*_CPTP–mem_〉 < −18 kcal/mol are labeled. The black line indicates the average position of phosphate groups of membrane lipids.

### CPTP alters the membrane’s physical properties to facilitate C1P extraction

When bound to the membrane, CPTP also influences the molecular structure and physical properties of the membrane. Previous simulations suggested that helix α6, in particular, can promote lipid reorientation into configurations apt for tail-first uptake by CPTP: By orienting lipids parallel with the membrane surface, the terminal carbons of their hydrophobic tails are positioned at the membrane surface and pointing into CPTP’s hydrophobic cavity [27]. Consistent with this idea, we find that the hydrophobic tails of lipids in close proximity to CPTP are oriented more parallel to the membrane surface than expected for a POPC membrane without CPTP bound (Fig 5). Lipid reorientation is roughly coupled to decreased lipid order (S5 Fig A-B) and membrane thickness (S5 Fig C-D). Furthermore, we find that not only helix *α*6, as suggested previously [27], but also the N-terminus of helix *α*2 can promote lipid reorientation (S6 Fig A). In its apo form, CPTP promotes increased lipid reorientation necessary for C1P uptake. Around the C1P-bound form of CPTP, reorientation of membrane lipids may enable the formation of initial hydrophobic contacts between C1P and the membrane, helping to guide C1P from CPTP’s hydrophobic cavity into the membrane. In fact, lipids in proximity to the C1P-bound form are typically oriented more parallel to the membrane surface and more disordered than those around the apo form (Fig 5B-C and S5 Fig A-B). We attribute this to differences in the conformations of the C1P-bound and apo forms (Fig 2, 3, and S1 Text, Note 1) that facilitate different types of interactions with membrane lipids. For example, in the C1P-bound form, helix *α*5, which stacks roughly on top of helix *α*6, is positioned such that nearby lipids exhibit increased reorientation and disorder compared to lipids in a POPC membrane without CPTP bound (S6 Fig A and B). In contrast, lipids in proximity to helix *α*5 of the apo form are oriented slightly more perpendicular to the membrane surface and increasingly ordered (S6 Fig A and B). A decrease in membrane order may, thus, greatly ease C1P insertion into a membrane.

**Fig 5.**
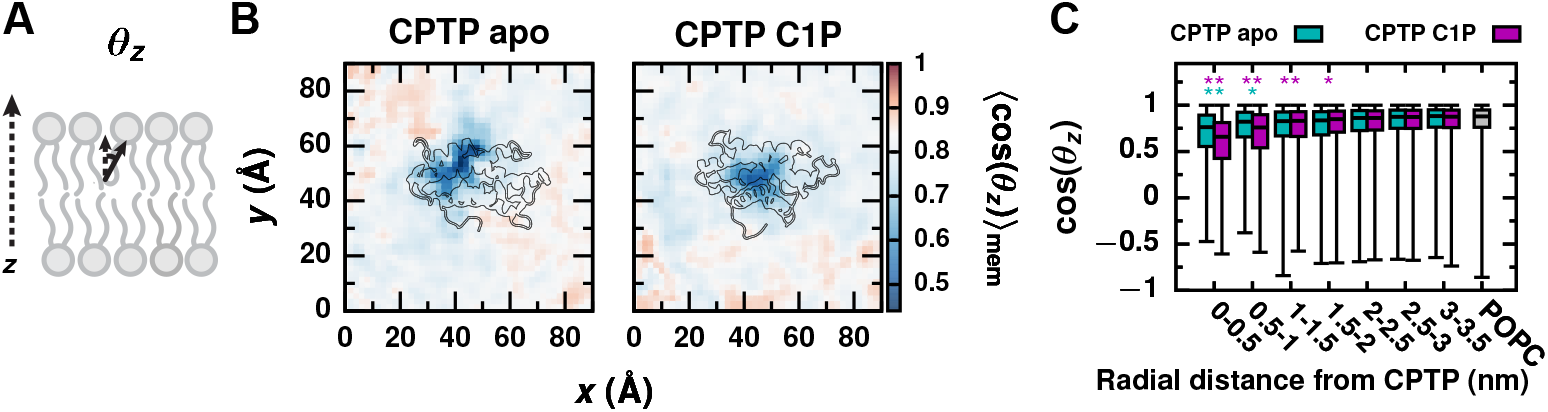
CPTP reorients nearby lipids in the membrane. (A) Lipid orientation is characterized by the angle *θ_z_* each lipid tail makes with the *z* axis, which is normal to the membrane surface. (B) Average cos(*θ_z_*) of membrane lipids as a function of a lipid’s displacement in the *xy* plane from CPTP. Color scale is set relative to 〈cos(θ_z_)〉_mem_ for a POPC membrane without CPTP present (white). Structures of the apo and C1P-bound forms of CPTP bound to the membrane are outlined. (C) Box-and-whisker plots of cos(*θ_z_*) for lipids within a specified radial distance from CPTP’s center-of-mass in the xy plane and for lipids in a POPC membrane without CPTP bound. The box extends from the 1st to 3rd quartile with the median indicated by the black line, and the whiskers extend from the minimum to maximum value. Asterisks indicate the average differs significantly from that of a POPC membrane: ** *p* < 0.001; * *p* < 0.01 (Welch’s t-test).

A membrane property that is especially key to lipid transport is the hydrophobicity of a lipid’s local membrane environment. We previously demonstrated that the activation free energy for passive lipid transport correlates with the average number of hydrophobic contacts a lipid makes with surrounding membrane lipids, 〈*n*_CC_〉_mem_ (Fig 6A) [26]. As 〈*n*_CC_〉_mem_ decreases, the rate of passive lipid desorption from a membrane increases [26]. We find that CPTP significantly reduces the number of hydrophobic contacts that nearby lipids make with other membrane lipids, and the apo form results in a greater disruption of a lipid’s local hydrophobic environment than the C1P-bound form. As shown in Fig 6B, a minimal 〈*n*_CC_〉_mem_ of 1,650 contacts (69% of the contacts formed on average between lipids in a membrane without CPTP bound) is observed near the entrance to the apo form’s hydrophobic cavity, whereas the minimal 〈*n*_CC_〉_mem_ around the C1P-bound form is 2,100 (88% of the contacts formed on average without CPTP bound). Fig 6C plots distributions of *n*_CC_ for membrane lipids as a function of radial distance from CPTP. Although radial averaging obscures geometric and anisotropic features evident in Fig 6B, the analysis in Fig 6C indicates that lipids within 5 Å of the apo form’s binding location make an average of 300 fewer hydrophobic contacts than lipids in a membrane without CPTP bound (*p* = 2.6 × 10^-9^, Welch’s t-test), whereas lipids within 5 Å of the C1P-bound form make an average of 250 fewer hydrophobic contacts (p = 4.9 × 10^-7^, Welch’s t-test). Based on the relationship between 〈*n*_CC_〉_mem_ and the free energy barrier for passive lipid desorption provided in [26], this reduction in 〈*n*_CC_〉_mem_ for lipids within 5 Å of the apo form corresponds to an expected decrease in the free energy barrier of 2 kcal/mol. Thus, CPTP may reduce the barrier for hydrophobic C1P–membrane contact breakage, which limits the rate of passive lipid transport [25, 26].

**Fig 6.**
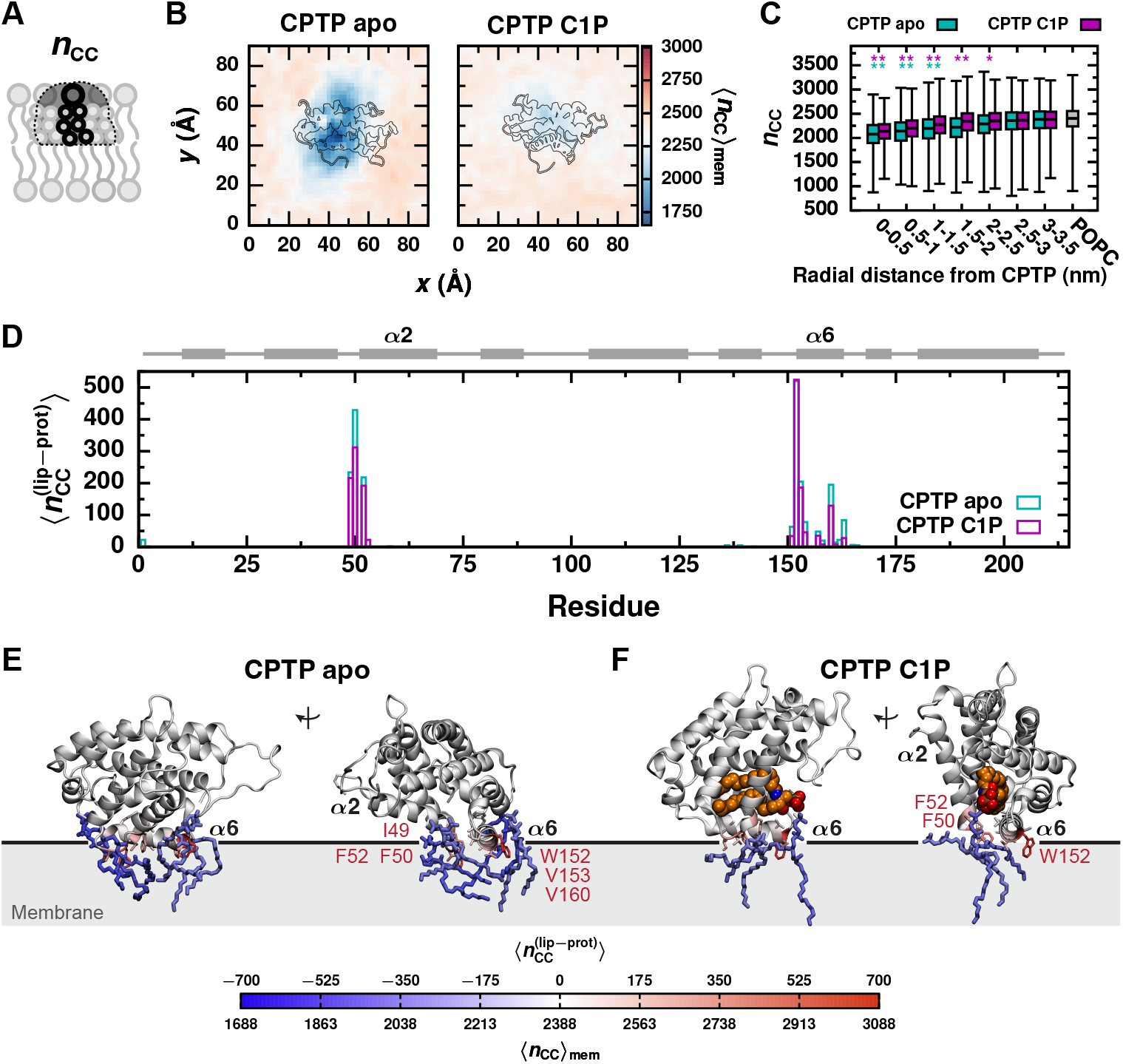
The apo form of CPTP significantly disrupts the local hydrophobic membrane environment of lipids below. (A) A lipid’s hydrophobic environment is characterized by the number of hydrophobic carbon–carbon contacts it makes with surrounding lipids, *n*_CC_. (B) Average *n*_CC_ of membrane lipids as a function of a lipid’s displacement in the *xy* plane from CPTP. Color scale is set relative to 〈*n*_CC_〉_mem_ for a POPC membrane without CPTP present (white). Structures of the apo and C1P-bound forms of CPTP bound to the membrane are outlined. (C) Box-and-whisker plots of ncc for lipids within a specified radial distance from CPTP’s center-of-mass in the *xy* plane and for lipids in a POPC membrane without CPTP bound. The box extends from the 1st to 3rd quartile with the median indicated by the black line, and the whiskers extend from the minimum to maximum value. Asterisks indicate the average differs significantly from that of a POPC membrane: ** p <0.001; * p <0.01 (Welch’s t-test). (D) Average number of hydrophobic contacts each residue of CPTP makes with membrane lipids, 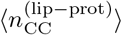. CPTP’s secondary structure is schematically illustrated above with helices represented as rectangles and unstructured loop regions as lines. (E and F) 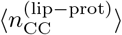 mapped onto the structures of the (E) apo and (F) C1P-bound forms. Residues with 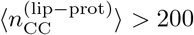 are labeled. Example configurations of lipids with (1) a reduction in *n*_CC_ of ≥ 300 contacts relative to 〈*n*_CC_〉_mem_ of a POPC membrane without CPTP bound and (2) ≥ 90% of those contacts replaced with hydrophobic contacts with CPTP are shown. Additional examples are shown in S7 Fig. The black line indicates the average position of phosphate groups of membrane lipids.

In general, decreases in 〈*n*_CC_〉_mem_ can arise from looser lipid packing, decreased membrane order, and membrane thinning [26]. We find that lipids around the apo form, exhibit decreased 〈*n*_CC_〉_mem_ largely due to looser lipid packing, as quantified by the area per lipid (S5 Fig E and F), as opposed to membrane thinning or decreased membrane order (S5 Fig A-D). Lipids with the largest reduction in *n*_CC_ and increase in occupied area are proximal to helices *α*1, *α*2, *α*5, and *α*6 (S6 Fig C and D). Hydrophobic residues on these helices can wedge themselves between lipid tails and, thereby, engage in hydrophobic lipid–protein contacts and alter lipid packing. In particular, residues F50 and F52 on helix α2 and W152 on helix *α*6 each engage in more than 200 hydrophobic lipid–protein contacts on average (Fig 6D). Indeed, these residues favorably interact with the membrane (Fig 4), to a large extent through van der Waals interactions. The thermodynamic cost of broken hydrophobic lipid–membrane contacts is thus mitigated by the formation of favorable lipid–protein interactions. Lipids with a reduced number of hydrophobic lipid–lipid contacts often from a compensatory number of hydrophobic contacts with CPTP; Fig 6E and 6F and S7 Fig show examples of such lipids. Because the apo form of CPTP binds deeper into the membrane (Fig 2), it forms more hydrophobic lipid–protein contacts than the C1P-bound form (Fig 6D) and breaks more hydrophobic lipid–membrane contacts (Fig 6B and C).

### CPTP rapidly extracts C1P by lowering the free energy barrier for breaking hydrophobic C1P–membrane contacts

To determine if CPTP lowers the activation free energy barrier for passive C1P desorption from a membrane, Δ*F*_des_ [25, 26], as hypothesized in the previous section, we performed all-atom free energy calculations for systems both with and without CPTP. Δ*F*_des_ is obtained from the free energy profile Δ*F* (*r*_LxS_) as a function of the reaction coordinate [32–34] for passive lipid (L) transport via solvent (xS), *r*_LxS_, defined in [26] and Eq 1 in the Methods. *r*_LxS_ measures hydrophobic lipid–membrane interactions and monitors progress from configurations with C1P in the membrane (which exhibit many hydrophobic C1P–membrane contacts and large positive values of *r*_LxS_) to configurations with C1P outside the membrane (which lack hydrophobic C1P–membrane contacts and have negative values of *r*_LxS_). ΔF_des_ is the largest free energy barrier along ΔF (*r*_LxS_) separating these two states (more precisely, 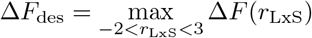 after setting Δ*F*(*r*_LxS_) = 0 at the global free energy minimum) [26].

However, because *r*_LxS_ only monitors C1P–membrane interactions, *r*_LxS_ alone cannot reliably identify if C1P is taken up by CPTP. So, for the system containing CPTP, we first calculate a 2D free energy surface, Δ*F*(*r*_LxS_, *Q*), as a function of *r*_LxS_ and a second order parameter *Q* that monitors C1P–CPTP interactions. More specifically, *Q* is the fraction of contacts C1P makes with CPTP when fully inside its hydrophobic cavity, including (a) hydrophobic contacts between C1P and residues lining CPTP’s cavity and (b) polar contacts between C1P’s headgroup and sphingoid backbone and residues responsible for C1P recognition. (A triad of cationic residues located within a positively-charged surface pocket at the entrance to the hydrophobic cavity engages C1P’s phosphate headgroup, and a bifurcated hydrogen bond recognizes C1P’s acyl-amide moiety [16].) Q distinguishes configurations with C1P outside CPTP (*Q* ≈ 0) from configurations with C1P fully housed within CPTP (*Q* > 0.8) (S1 Text, Note 2), and is calculated using the functional form developed to calculate the fraction of native protein contacts [35] and defined in Eq 4 in the Methods. We then numerically integrate Δ*F* (*r*_LxS_, *Q*) over *Q* to obtain the free energy profile Δ*F* (*r*_LxS_) and determine Δ*F*_des_ in the presence of CPTP. To efficiently calculate Δ*F*(*r*_LxS_, *Q*), we performed multi-walker [36] well-tempered metadynamics simulations [37] biasing *r*_LxS_ and *Q* to enhance the sampling of C1P–membrane and C1P–CPTP interactions, respectively. Initiating walkers from both apo and C1P-bound forms of CPTP bound to the membrane further facilitated sampling of both forward and reverse transitions between configurations with C1P in the membrane and configurations with C1P inside CPTP, and convergence was obtained after 500 ns (cumulatively 10 *μ*s) (S8 Fig).

Results of the free energy calculations are shown in Fig 7. At the global free energy minimum of Δ*F*(*r*_LxS_, *Q*) (outlined in cyan in Fig 7A), C1P resides in the membrane where it makes many hydrophobic contacts with surrounding membrane lipids but does not substantially interact with CPTP. An example of a configuration found within the global free energy minimum is shown in Fig 7C1. At the second most favorable minimum of Δ*F*(*r*_LxS_, *Q*) (outlined in magenta in Fig 7A), C1P resides within CPTP’s hydrophobic cavity but does not form hydrophobic contacts with membrane lipids. An example of a configuration found at this local minimum is shown in Fig 7C12. The free energy surface Δ*F*(*r*_LxS_, *Q*) is consistent with CPTP’s observed function as a C1P transporter: CPTP serves as a free energetically favorable vehicle for C1P transport between even more favorable states in which C1P is part of a membrane.

**Fig 7.**
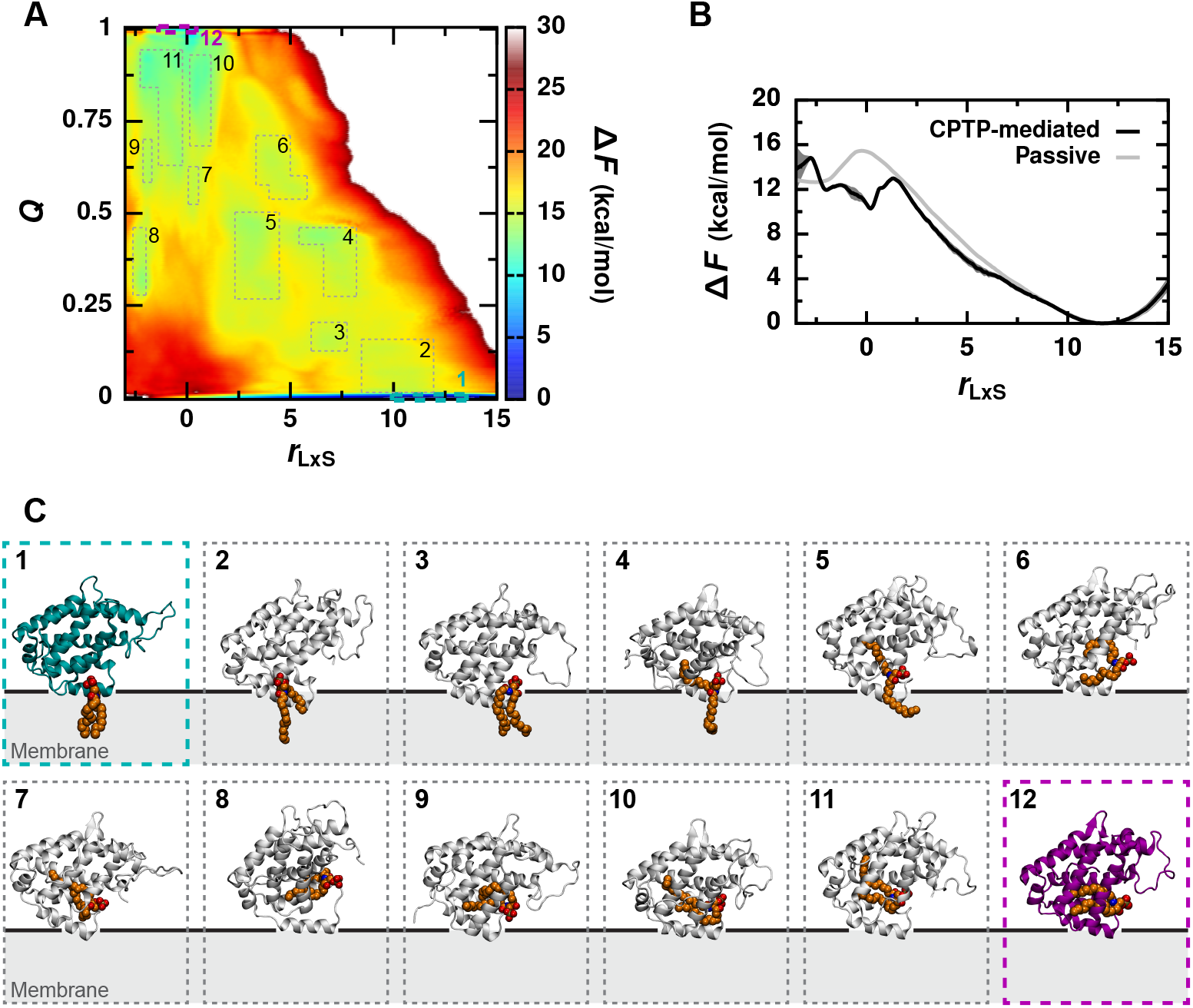
CPTP lowers the free energy barrier for hydrophobic C1P–membrane contact breakage. (A) Free energy surface Δ*F*(*r*_LxS_,*Q*). Error is shown in S9 Fig. (B) Free energy profile Δ*F*(*r*_LxS_) for CPTP-mediated C1P extraction and passive C1P desorption. For CPTP-mediated extraction, Δ*F*(*r*_LxS_) is obtained by marginalizing Δ*F*(*r*_LxS_, *Q*) over *Q*, and error bars indicate standard error computed with block averaging. (Δ*F*(*Q*) obtained analogously is shown in S10 Fig.) For passive C1P desorption, Δ*F*(*r*_LxS_) is computed with umbrella sampling simulations, convergence is demonstrated in S11 Fig, and the error is comparable to the line width. (C) Example configurations from different regions of the free energy surface outlined in (A). C1P is rendered as van der Waals spheres and colored orange. The apo form of CPTP in configuration 1 is colored dark cyan, and the C1P-bound form in configuration 12 is colored dark magenta. The black line indicates the average position of phosphate groups of membrane lipids. Corresponding views looking into the hydrophobic cavity are shown in S12 Fig.

Fig 7B shows Δ*F*(*r*_LxS_) both in the presence and absence of CPTP. The free energy profiles match at large values of *r*_LxS_ as expected—being a catalyst, CPTP does not alter the relative probability of configurations within the global free energy minimum around *r*_LxS_ ≈11.8. Passive C1P transport has a rate-limiting activation free energy barrier of Δ*F*_des_= 15.46 ±0.08 kcal/mol at *r*_LxS_ ≈0. When CPTP is present, Δ*F*_des_ = 13.0 ±0.2 kcal/mol, and the barrier is located at slightly larger values of *r*_LxS_. CPTP lowers Δ*F*_des_by 2.5 ±0.2 kcal/mol, in agreement with the value predicted in the previous section based on the average reduction in 〈*n*_CC_〉_mem_ caused by the apo form. By disrupting C1P’s local hydrophobic environment in the membrane (Fig 6), CPTP lowers the free energy barrier for hydrophobic lipid–membrane contact breakage and, thus, catalyzes lipid extraction. Because CPTP provides C1P with another favorable hydrophobic environment, C1P enters CPTP’s hydrophobic cavity after fewer hydrophobic contacts with the membrane have been broken than as required for C1P desorption into solvent, which offers no favorable hydrophobic environment. As a result, the free energy barrier Δ*F*_des_ occurs at slightly larger values of *r*_LxS_ when CPTP is present. Indeed, as illustrated by an example configuration near this free energy barrier in Fig 7C5, C1P maintains hydrophobic contacts with the membrane while also forming contacts with CPTP.

The free energy calculations (Fig 7) suggest that C1P entry into CPTP’s hydrophobic cavity may require passage through multiple metastable intermediates. Configurations representative of potential intermediates are shown in Fig 7C. Configurations 2 and 3 shown in Fig 7C illustrate how interactions with helices *α*6 and *α*2 can position C1P for tail-first entry into CPTP’s hydrophobic cavity. Based on the observation of two C1P-binding modes in crystal structures, C1P entry and exit have been suggested to occur through a two-step mechanism whereby the acyl chain enters into the cavity before the sphingosine during entry and leaves after the sphingosine during exit [16]. Consistent with this, we observe configurations with one tail of C1P inside CPTP’s hydrophobic cavity and the other still within the membrane (configurations 4 and 5 in Fig 7C). We do not observe a preference for one tail over the other to be housed within the cavity. For example, the acyl chain is found in the cavity in configurations 5 and 6 shown in Fig 7C, whereas the sphingosine chain is found in the cavity in configurations 2, 4, and 8. Nevertheless, our findings support a two-step mechanism whereby each tail of C1P enters into and exits from CPTP’s hydrophobic cavity individually. Once C1P has fully broken hydrophobic contacts with the membrane, it can adopt various configurations inside CPTP’s hydrophobic cavity as illustrated by configurations 9-12 in Fig 7C. Collectively, the intermediates observed in our simulation suggest that CPTP may extract C1P through multiple, molecularly diverse mechanisms.

## Discussion

Cytosolic shuttle-like lipid transfer proteins enable rapid alteration of membrane compositions through trafficking of individual lipids [10–14]. Yet, how they swiftly extract individual lipids from a membrane remains an outstanding question. Our recent discovery that the rate of passive lipid transport is limited by the disruption of a lipid’s local hydrophobic environment [25, 26] suggests that lipid transfer proteins may catalyze lipid extraction by lowering the free energy barrier for hydrophobic lipid–membrane contact breakage. Here, we tested this hypothesis for the specific case of CPTP-mediated transport of the bioactive sphingolipid C1P. To do so, we devised a consecutive multiscale simulation scheme tailored to studying lipid transport via lipid transfer proteins: We used coarse-grained MD simulations to observe multiple spontaneous membrane binding events, which occur over microsecond timescales, and then used all-atom MD simulations to resolve key conformational changes that facilitate C1P uptake and to quantify CPTP’s ability to catalyze C1P extraction from a membrane. Such consecutive multiscale schemes are ideal for studying process whose elementary steps occur at varied time and length scales, such as lipid transfer cycles. Indeed, a similar approach was recently used to investigate influenza hemagglutinin driven membrane fusion [38], a process related to vesicular lipid transport [12, 39].

Based on our simulations, we propose a molecular mechanism for how CPTP functions as a cytosolic C1P shuttle that is consistent with the previously suggested cleft-like gating mechanism [16]: To initiate a transfer event, the apo form of CPTP, whose collapsed hydrophobic cavity is closed off to C1P entry, binds a donor membrane. Interactions between the membrane and helix *α*6 in addition to gating helix *α*2 anchor CPTP deeply into the membrane and aptly position the entrance to its hydrophobic cavity at the membrane–solvent interface. Strong electrostatic interactions between positively charged residues on gating helix *α*2 may specifically facilitate cleft widening and gate opening. Poised for C1P entry into its hydrophobic cavity, CPTP disrupts C1P’s local hydrophobic membrane environment by engaging in hydrophobic lipid–protein contacts. This lowers the free energy barrier for passive C1P desorption from the membrane and enables CPTP to catalyze C1P extraction. Additionally, interactions between C1P and residues on helices *α*2 and *α*6 orient C1P for tail-first uptake into CPTP’s hydrophobic cavity. Entry proceeds through a two-step mechanism in which one tail of C1P enters prior to the other. Complete uptake of C1P into CPTP’s hydrophobic cavity further opens gating helix *α*2. Such conformational changes may help break strong electrostatic interactions with the membrane that had previously anchored the apo form and, thus, potentially facilitate membrane unbinding. Having extracted C1P from a donor membrane, CPTP then inserts C1P into an acceptor membrane, a free energetically downhill event, to complete a transfer cycle.

Based on the reduction in the free energy barrier for hydrophobic lipid–membrane contact breakage calculated in our simulations, we estimate that CPTP-mediated extraction of C1P is roughly two orders-of-magnitude faster than passive C1P desorption. In *in vitro* lipid transfer assays, CPTP transfers approximately 4 C1P molecules per minute [16, 19, 20]. While the rate of passive C1P transport is expected to be at least two orders-of-magnitude slower [19–24], to our knowledge, it has not yet been measured experimentally, hindering a quantitative comparison between our simulations and experiment. As determined in our previous studies [25, 26], both simulation and experiment yield the same values for changes in the activation barrier for passive lipid transport, for example due to differences in membrane physicochemical properties, even though the absolute barrier height is severely underestimated in simulation. This suggests that simulations provide accurate physical insights into lipid transport but lack quantitative accuracy, for example, due to limitations in additive force fields [40–42]. Therefore, while the reduction in the free energy barrier for hydrophobic contact breakage by CPTP calculated in our simulations may prove to match experimental values, the absolute barrier heights reported herein will almost certainly differ from experimental ones.

Nevertheless, the mechanism proposed herein offers new insights into the role of hydrophobic residues that are conserved between CPTP and the *Arabidopsis* C1P transfer protein, ACD11 [43], and shared with other members of the GLTP family [18, 28–30]. We identify F50, F52, and W152 as important for enhancing C1P extraction from the membrane: By forming hydrophobic contacts with lipid tails, these residues create a wedge between lipids and break hydrophobic lipid–membrane contacts. Of these residues, F50 and W152 engage in the largest number of hydrophobic lipid–protein interactions, a potential biophysical reason for their high conservation among GLTP family members [18, 28–30, 43]. In ACD11, phenylalanine residues analogous to F50 and F52 on helix *α*2 of CPTP serve as “portal gates” that swing open to allow C1P entry into it’s hydrophobic cavity [43]. Our results suggest that swinging open of these “portal gates” not only allows for C1P entry but, in fact, facilitates it by disrupting C1P’s local hydrophobic environment in the membrane. In this way, helix *α*2 plays dual roles as hydrophobic cavity gate and membrane disruptor.

Our proposed mechanism also offers new hypotheses for how certain anionic lipids may enhance C1P transfer by CPTP. Including PS, PI(4,5)P_2_, or PI4P, but not other anionic lipids, in donor membranes has been shown to enhance C1P transport in in *vitro* assays [19, 20]. Through specific interactions with CPTP’s helix *α*6 and *α*3-*α*4 loop, these lipids help position the entrance to CPTP’s hydrophobic cavity at the membrane surface and regulate its activity [19, 20, 27]. We hypothesize that PS, PI(4,5)P_2_, and PI4P may also interact with arginine and lysine residues on helix *α*2 when CPTP is bound to the membrane in its apo form. Compared to other anionic lipids, PS, PI(4,5)P_2_, and PI4P have larger headgroups with terminal acidic groups that may allow them to interact with these basic residues on helix *α*2, which are located farther above the membrane surface than residues on helix *α*6. Such interactions may facilitate binding of the apo form to the membrane and also promote opening of gating helix *α*2 and widening of CPTP’s hydrophobic cavity as necessary for C1P uptake. Systematically varying membrane composition in simulations using the multiscale approach presented here offers an avenue to test these hypotheses. Such studies are key to ultimately understanding how complex cell membrane compositions [44–46] impact CPTP-mediated C1P transfer and how CPTP maintains net flow of C1P from the Golgi to plasma membrane [16].

Our proposed mechanism for C1P transfer via CPTP additionally shares biophysical features with mechanisms of other cytosolic shuttle-like lipid transfer proteins, despite differences in their structures and lipid specificities. For example, members of the StARkin (relatives (kin) of steroidogenic acute regulatory protein (StAR)) superfamily, which have *α*-*β* topologies and bind phospholipids in the opposite orientation to CPTP (head-first versus tail-first) [14, 47], also undergo conformational changes upon membrane binding that facilitate opening of their hydrophobic cavities [48–52]. Lipid transfer proteins in the Sec14 and oxysterol-binding protein (OSBP)-related protein (ORP) families, which have well-defined lids to their hydrophobic cavities and bind phospholipids tail-first [14, 47], similarly exhibit conformational changes upon membrane binding that displace the lids to their hydrophobic cavities as required for lipid uptake [53–55]. Thus, our results add to the growing evidence that lipid transfer proteins couple conformational changes to membrane binding to regulate lipid uptake and release.

Electrostatic interactions between basic protein residues and anionic lipids can promote the aforementioned conformational changes and aid lipid uptake [49, 53, 54]. Similarly, we identify a potential role for electrostatic interactions in facilitating opening of CPTP’s gating helix *α*2. Complete opening of this gate to accommodate C1P inside CPTP’s hydrophobic cavity breaks these favorable electrostatic interactions. We suggest this may enable membrane unbinding in a manner reminiscent of the electrostatic switching mechanism used by Osh6p, a member of the ORP family that has a well-defined lid to its hydrophobic cavity [55]. Such switching mechanisms, in which a lipid transfer protein’s membrane affinity depends on lipid occupancy, enable efficient transfer cycles by ensuring membrane binding is transient [55, 56].

Significantly, we demonstrate how CPTP enhances C1P extraction by breaking hydrophobic C1P–membrane contacts and lowering the activation free energy barrier for passive lipid desorption. To our knowledge, this offers the first molecular evidence that a lipid transfer protein catalyzes lipid extraction from a membrane. Because the catalytic mechanism described here is closely related to that of passive lipid transport [25, 26], we suspect this biophysical mechanism may be generally used by lipid transfer proteins, not just CPTP, to increase the rate of lipid transport between cell membranes. For example, the sterol transfer protein StarD4 has been shown to deform donor membranes [57], as one would expect for a protein that catalyzes lipid extraction by disrupting a lipid’s local hydrophobic environment [26]. Nevertheless, we expect the precise molecular details to vary from protein to protein. Our mulitscale simulation approach provides a framework to test this general hypothesis and illuminate the variety of molecular mechanisms employed by lipid transfer proteins to alter a lipid’s membrane environment and catalyze its extraction. Doing so will advance our understanding of how lipid transfer proteins rapidly traffic lipids to ensure membrane homeostasis.

## Methods

### Molecular dynamics simulations

We performed molecular dynamics simulations of CPTP both in solution and in the presence of a POPC membrane at both all-atom and coarse-grained resolutions using GROMACS 2019 [58]. Simulations were conducted in 150 nM NaCl solutions, matching the conditions used in *in vitro* experiments of C1P transfer by CPTP [16, 19, 20]. The CHARMM36 force field [59] in combination with the CHARMM TIP3P water model [60] was used for the all-atom simulations. CHARMM36 force field parameters developed for sphingolipids [61] were used for C1P with a saturated acyl chain of 16 carbons (16:0-C1P). The MARTINI 2.3P force field [62] in combination with polarizable MARTINI water model [63] was used for all coarse-grained simulations since it accurately reproduces the binding poses of peripheral membrane proteins [62]. An ElNeDyn elastic network [64] with harmonic springs connecting all backbone beads within 0.9 nm of each other and a spring constant of 500 kJ/mol/nm^2^ was used to maintain the overall shape of CPTP in its apo and C1P-bound forms. Parameters for 16:0-C1P were chosen according to the standard MARTINI 2.0 lipid definitions and building block rules [65].

#### All-atom solution-phase simulations

The crystal structure of CPTP provided in PDB 4K85 [16] was used to initialize all-atom simulations since it has the most residues fully resolved out of all structures of CPTP with C1P fully bound. The first six N-terminal residues that were not resolved in PDB 4K85 were built using MODELLER [66, 67]. To investigate the transfer of CPTP’s most likely *in vivo* substrate, the structure of 16:0-C1P fully bound inside CPTP provided in PDB 4K84 was aligned onto the structure of C1P with a saturated acyl chain of 12 carbons (12:0-C1P) present in PDB 4K85 to generate an initial configuration of CPTP in its C1P-bound form. C1P was removed from this structure to generate an initial configuration of the apo form. Both initial structures of the apo and C1P-bound forms were solvated in triclinic boxes of pre-equilibrated water molecules with dimensions chosen such that the minimum distance between any protein atom and the box walls was at least 2 nm. Sodium and chloride ions were added to achieve a salt concentration of 150 mM and to fully neutralize each system. Both systems were then energy minimized using the steepest descent algorithm and equilibirated using a three-step protocol: (1) a 200 ps simulation in an isothermal–isochoric (NVT) ensemble with the positions of CPTP’s heavy atoms restrained by harmonic potentials with a force constant of 1000 kJ/mol/nm^2^ to equilibrate the solvent molecules and adjust the system density; (2) a 200 ps simulation in an isothermal–isobaric (NPT) ensemble using the Berendsen barostat [68] to maintain the pressure with isotropic pressure coupling and a coupling time constant of 2 ps; and (3) a 200 ps simulation in an NPT ensemble using the Parinello-Rahman barostat [69] with a coupling time constant of 5 ps.

Subsequently, production simulations in an NPT ensemble were run for 300 ns using the parameters given below. The final 200 ns of each of these simulations was analyzed to characterize the solution-phase structure of the apo and C1P-bound forms of CPTP. The final configurations from these simulations were used to initialize simulations with a POPC bilayer present.

#### Coarse-grained simulations of CPTP in the presence of a membrane

Coarse-grained bilayers of 264 lipids surrounded by 3.2 nm slabs of solvent were built using INSANE [70] for simulations of CPTP in the presence of a membrane. For simulations of the apo form, the bilayer consisted of 262 POPC lipids and two 16:0-C1P lipids, one in each leaflet. For simulations of the C1P-bound form, the bilayer consisted of 264 POPC lipids. Prior to the addition of CPTP into the solvent surrounding the bilayer, the structures of both bilayers were fully relaxed from their initial lattice configurations. This involved an energy minimization using the steepest descent algorithm followed by equilibration using a two-step protocol: (1) a 500 ps simulation using a 10 fs time step and the Berendsen barostat [68] with semi-isotropic presssure coupling and a coupling time constant of 3 ps; and (2) a 1 ns simulation using a 20 ps time step and the Parinello-Rahman barostat [69] with a coupling time constant of 12 ps. Subsequently, a 50 ns production run was performed to obtain bilayer structures that were used to construct initial configurations of CPTP in the presence of a membrane.

The martinize.py script [71] was used to construct coarse-grained protein topologies and structures based on the final configurations of the apo and C1P-bound forms of CPTP obtained from all-atom solution-phase simulations. Both coarse-grained structures were then fully equilibrated in solution through an energy minimization and a three-step protocol: (1) a 500 ps simulation in an NVT ensemble using a 10 fs time step and with the positions of CPTP’s backbone beads restrained by harmonic potentials with a force constant of 1000 kJ/mol/nm^2^ to equilibrate solvent molecules and adjust the system density; (2) a 500 ps simulation in an NPT ensemble using a 20 fs time step and the Berendsen barostat [68] with isotropic pressure coupling and a coupling time constant of 3 ps; and (3) a 500 ps simulation using a 20 fs time step and the Parinello-Rahman barostat [69] with a coupling time constant of 12 ps. Subsequently, 300 ns production runs were performed to obtain structures of CPTP used to initialize simulations that probe membrane binding.

For simulations of both forms of CPTP in the presence of a membrane, four different initial coarse-grained configurations were built by inserting a structure of CPTP sampled during the last 100 ns of the coarse-grained solution-phase simulation in a random location and orientation in the solvent around the equilibrated bilayer such that the minimum distance between any atom of CPTP and the membrane was 6 Å. Each replicate was then energy minimized and equilibrated using a three-step protocol: (1) a 500 ps simulation in an NVT ensemble using a 10 fs time step and with the positions of CPTP’s backbone beads restrained by harmonic potentials with a force constant of 1000 kJ/mol/nm^2^ to equilibrate solvent molecules and adjust the system density; (2) a 500 ps simulation in an NPT ensemble using a 20 fs time step and the Berendsen barostat [68] with semi-isotropic pressure coupling and a coupling time constant of 3 ps; and (3) a 1 ns simulation using a 20 fs time step and the Parinello-Rahman barostat [69] with a coupling time constant of 12 ps. During production runs of 2 *μ*s, CPTP spontaneously bound to the membrane in all but one replicate of the apo form (S2 Fig), which was excluded from analysis of CPTP’s membrane binding pose in coarse-grained simulations.

#### All-atom simulations of CPTP bound to a membrane

Membrane-bound coarse-grained configurations of the apo and C1P-bound forms of CPTP were then backmapped into all-atom representations using the backward method [72]. The systems then underwent a three-step equilibration using the same protocol as used for the all-atom solution-phase simulations but with semi-isotropic pressure coupling. Subsequently, 300 ns production runs were performed. The final 200 ns from each of these runs was used to analyze all-atom configurations of CPTP bound to a membrane.

#### All-atom production simulation parameters

All simulations were performed in an NPT ensemble at a pressure of 1 bar and temperature of 310 K. Separate Nosé-Hoover thermostats [73, 74] were used for the solvent and solute molecules. The pressure was maintained using the Parinello-Rahman barostat [69] with a compressibility of 4.5 ×10^−5^ bar^−1^ and coupling time constant of 1 ps. Isotropic pressure coupling was used in solution-phase simulations, and semi-isotropic coupling was used in simulations with a bilayer. Dynamics were evolved using the leapfrog algorithm [75] and a 2 fs time step. All bonds to hydrogen were constrained using the LINCS algorithm [76]. Lennard-Jones forces were smoothly switched off between 0.8 and 1.2 nm. Real-space Coulomb interactions were truncated at 1.2 nm, and long-ranged Coulomb interactions were calculated using Particle Mesh Ewald (PME) summation [77]. Neighbor lists were constructed using the Verlet list cut-off scheme [78].

#### Coarse-grained production simulation parameters

All simulations were performed in an NPT ensemble at a pressure of 1 bar and temperature of 310 K. To avoid the “hot solvent–cold solvent” problem [79], separate V-rescale thermostats [80] were used for the solvent and solute molecules. The pressure was maintained using the Parinello-Rahman barostat [69] with a compressibility of 3 ×10^−4^ bar^−1^ and coupling time constant of 12 ps. Isotropic pressure coupling was used in solution-phase simulations, and semi-isotropic coupling was used in simulations with a bilayer. Dynamics were evolved using the leapfrog algorithm [75] and a 20 fs time step. Lennard-Jones and Coulomb interactions were truncated at 1.1 nm, and long-ranged Coulomb interactions were calculated using PME summation [77]. A relative dielectric constant of 2.5 was used. Neighbor lists were constructed using the Verlet list cut-off scheme [78].

### Analysis of CPTP’s structure and interactions with the membrane

We analyzed the structure of apo and C1P-bound forms of CPTP both in solution and bound to a membrane. Analysis was performed using a combination of MDAnalysis [81] and NumPy [82] Python libraries in addition to GROMACS tools [58] and MDpocket [31].

#### Hydrophobic cavity volume

We quantified the volume of CPTP’s hydrophobic cavity using MDpocket [31]. Only protein atoms were considered for this analysis. First, simulation frames were aligned to a common reference, specifically the C*α* atoms of residues 10 – 208 of the last frame of the solution-phase simulation of the C1P-bound form of C1P. This selection of residues excludes the flexible, unstructured termini. Then, the region at the interior of CPTP unoccupied by protein heavy atoms during at least 20% of the solution-phase simulation of the C1P-bound form was identified for analysis. The volume unoccupied in this region during solution-phase simulations of both the apo and C1P-bound forms was used as a measure of hydrophobic cavity volume.

#### Insertion depth

The insertion depth of each residue was calculated as the average signed distance in *z*, which is the axis normal to the membrane, between the center of mass (COM) of each residue and the average position of phosphate atoms (beads) in all-atom (coarse-grained) simulations.

#### Orientation of gating helix *α*2

To calculate the angles *θ*_*α*2_ and *ϕ*_*α*2_ (Fig 3B-C), the C*α* atoms of residues 54 and 65 were used to define the vector along the axis of helix *α*2, and an internal coordinate system was defined by the plane containing helix *α*6. The C*α* atoms of residues 155, 158, and 162 were used to define the plane that contains the top surface of helix *α*6 when CPTP is membrane bound. *θ*_*α*2_ and *ϕ*_*α*2_ are the polar and azimuthal angles, respectively, of the *α*2 vector in this internal coordinate system. *ϕ*_*α*2_ was specifically calculated as the angle between the projection of the *α*2 vector onto the plane and the vector along the axis of helix *α*6 defined by the C*α* atoms of residues 155 and 162.

#### CPTP–membrane interaction energy

The interaction energy between each residue of CPTP and the membrane, *E*_CPTP–mem_, was calculated as the sum of short-ranged Lennard-Jones and Coulomb interaction energy terms between each residue and the membrane.

#### Lipid tail orientation

The orientation of each lipid tail was quantified by the angle *θ_z_* (Fig 5A) between the z axis, which is normal to the membrane surface on average, and the vector connecting the terminal carbon of each tail to atom C2 for POPC or to atom C2S for C1P.

#### Number of hydrophobic contacts between lipids

The number of hydrophobic contacts between lipids, *n*_CC_, (Fig 6A) was calculated based on the distances 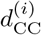 between hydrophobic carbons of two different lipids, where i labels a particular carbon-carbon (CC) pair. For all-atom POPC lipids, hydrophobic carbons include atoms C23-C216 and C33-C316. For all-atom C1P, hydrophobic carbons include atoms C3F-C16F and C5S-C18S. *n*_CC_ is the number of CC pairs that satisfy 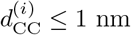.

#### Number of hydrophobic contacts between CPTP and lipids

The number of hydrophobic contacts formed between lipids and protein residues, 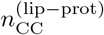, was calculated analogously to *n*_CC_. Specifically, 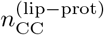 was calculated based on the distances 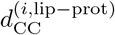 between a hydrophobic carbon of a lipid and a hydrophobic carbon of CPTP, where i labels a particular carbon-carbon (CC) pair. For all-atom CPTP, hydrophobic carbons include side chain carbons of Ala, Ile, Leu, Met, Phe, Trp, Val, Pro, and Cys residues. For all-atom lipids, hydrophobic carbons are defined as for *n*_CC_. 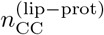 is the number of lipid–protein CC pairs that satisfy 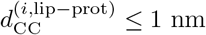.

#### Area per lipid

The area per lipid, *A*_lip_, was estimated as the area of the Voronoi cell containing each lipid calculated with FATSLiM [83] and using the phosphorous atom to define each lipid’s headgroup position. Because CPTP is only peripherally bound to the membrane, no protein atoms were taken into account to calculate *A*_lip_.

#### Membrane thickness

The membrane thickness was calculated as the average distance in z between the phosphorous atoms of lipids in the top and bottom leaflets.

#### Lipid acyl chain order parameter

Lipid acyl chain order was characterized by the average deuterium order parameter of all carbons in a lipid’s tails, *S*_CC_, as used to quantify membrane order in Seo et al. [84]. *S*_CC_ was calculated using LiPyphilic [85].

#### Significance testing

To determine if average membrane properties in the vicinity of CPTP differ significantly from those of a POPC membrane without CPTP bound, we obtained uncorrelated samples from our simulations using the method of Flyvbjerg and Petersen [86], and used those samples to compute p-values using Welch’s t-test [87].

### Free energy calculations

#### CPTP catalyzed C1P transport

We calculated free energy surfaces along the reaction coordinate for passive lipid transport, *r*_LxS_, and fraction of contacts between C1P and CPTP, *Q*. As determined in [25], *r*_LxS_ is a linear combination of two order parameters that measure hydrophobic lipid–membrane contacts between the transferring lipid and closest membrane leaflet. Both parameters are based on the collection of distances 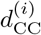 between a hydrophobic carbon of the lipid and a hydrophobic carbon of the membrane, where *i* labels a particular carbon-carbon (CC) pair: (1) 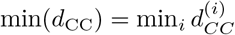 is the minimum of these CC distances; and (2) *n*_cc_ is the number of CC pairs that satisfy 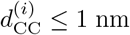. In detail,

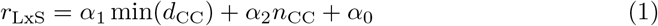

with coefficients *α*_1_ = −2.247 nm^-1^, *α*_2_ = 0.004828, and *α*_0_ = 0.6014 such that *r*_LxS_ is a unitless quantity [26]. During all enhanced sampling simulations, *r*_LxS_ was calculated using a differentiable form for min(*d*_CC_),

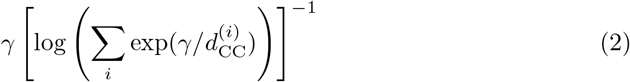

with *γ* = 200 nm, and for *n*_CC_,

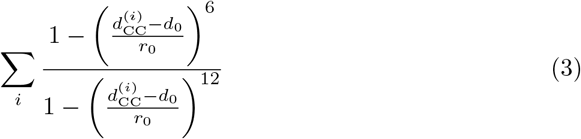

with *d*_0_ = 1 nm and *r*_0_ = 0.025 nm. *Q* is calculated using the functional form developed to calculate the fraction of native protein contacts [35], specifically,

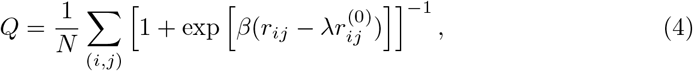

where *N* is the total number of contact pairs (*i, j*) between an atom *i* of C1P and atom *j* of CPTP, *r_ij_* is the distance between atoms *i* and *j*, 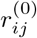 is a reference distance between atoms *i* and *j*, which we take to be the average distance between the atom pair in solution-phase simulations of the C1P-bound form, *β* = 50 nm^-1^, and *λ* = 1.8.

Contact pairs used to calculate *Q* include: (1) pairs of hydrophobic carbons of C1P and carbons of the residues that line CPTP’s hydrophobic cavity that have 〈*r_ij_*〉 ≤7.8 A during solution-phase simulations of the C1P-bound form of CPTP (residues 10, 14, 36, 39, 40, 42, 43, 46, 48, 50, 52, 53, 57, 110, 111, 114, 117, 118, 121, 122, 146, 150, 153, 154, 158, 162, 165, 171, and 175); and (2) pairs of heavy atoms of C1P’s headgroup and sphingoid backbone and heavy atoms of the residues responsible for C1P recognition that have have 〈r_ij_〉 ≤ 5.5 A during solution-phase simulations of the C1P-bound form (residues 52, 53, 56, 60, 96–102, 106, 110, 113, 114, 146–151, 154, and 214). Further discussion of the choice of contact pairs used to define Q is provided in Supporting Note 2 of S1 Text.

We used multi-walker [36] well-tempered metadynamics simulations [37] to calculate the free energy of CPTP-mediated C1P transport whereas, as described below, we used umbrella sampling simulations to calculate that of passive C1P transport. Metadynamics simulations can be more computationally efficient than umbrella sampling simulations [88]—a potential advantage when calculating free energy profiles for the larger system with CPTP present. Since both methods yield the same free energy estimates when converged [88], the choice of method is merely one of computational expense. Using multiple interacting walkers enables further computational gains through parallelization [36], and has been successfully used to efficiently simulate, for example, antibiotic translocation across membranes via porins [89]. To perform the multi-walker well-tempered metadynamics simulation we used the PLUMED 2 patch [90] for GROMACS. A total of 20 walkers, 10 initialized from all-atom configurations of the apo form of CPTP bound to the membrane and 10 initialized from all-atom configurations of the C1P-bound form bound to the membrane, were used. Gaussian hills with *σ*_*r*_LxS__ = 0.1 and *σ_Q_* = 0.01 and initial heights of 2.5 kJ/mol were added by each walker every 4 ps with a bias factor of 30. To avoid pushing the system into regions with very negative values of *r*_LxS_, where C1P is free in solution far away from either the membrane or CPTP, and at very large positive values of *r*_LxS_, which are unphysical, harmonic wall potentials with a spring constant of 500 kJ/mol were applied at *r*_LxS_ = −3 and 15. The simulation was run for 560 ns. Then, to facilitate sampling of configurations with C1P inside CPTP by walkers initialized with the apo form, and vice versa for walkers initialized with the C1P-bound form, moving restraints were applied to each walker to pull it to (*r*_LxS_ = −2.5, *Q* = 1), if it was initialized from the apo form, and to (*r*_LxS_ = 12, *Q* = 0), if it was initialized from the C1P-bound form. The moving restraint applied a harmonic bias on *r*_LxS_ with a spring constant of 75 kJ/mol and a harmonic bias on Q with a spring constant of 7500 kJ/mol over the course of 20 ns. During this time, no Gaussians were deposited, but the bias due to previously accumulated Gaussians was applied. Subsequently, the multi-walker well-tempered metadynamics simulation was continued for an additional 560 ns without the moving restraints. In total, the simulation was run for 1.14 *μ*s (cumulatively 22.8 *μ*s). Convergence was obtained after 500 ns (S8 Fig). After discarding the data from the first 500 ns of each walker and from the time period when moving restraints were applied, a 2D free energy surface Δ*F*(*r*_LxS_, *Q*) and 1D free energy profiles Δ*F*(*r*_LxS_) and Δ*F*(*Q*) were obtained using the time-independent free energy estimator of Tiwary and Parrinello [91]. This amounts to marginalizing Δ*F*(*r*_LxS_,*Q*) over one variable to obtain the 1D free energy profiles Δ*F*(*r*_LxS_) and Δ*F*(*Q*). Error bars were calculated as the standard error of the free energy estimated from two independent 310 ns blocks.

#### Passive C1P transport

We calculated a free energy profile along *r*_LxS_ for passive transport of C1P between membranes using umbrella sampling simulations [92] as done in our previous study [26]. Simulations were performed using two different membrane compositions that matched the membrane compositions used for simulations with both forms of CPTP. An initial bilayer of 264 POPC lipids surrounded by 3.2 nm thick slabs of solvent was built using the CHARMM-GUI Membrane Builder [93, 94]. A lipid in each leaflet was replaced with a 16:0-C1P molecule to construct a second membrane. As in the simulations of CPTP in the presence of a membrane, sodium and chloride ions were added to achieve a salt concentration of 150 mM and to fully neutralize each system. Both membranes were energy minimized and equilibrated using a two-step protocol: (1) a 250 ps simulation using the Berendsen barostat [68]; and (2) a 250 ps simulation using the Parinello-Rahman barostat [69]. All production simulation parameters matched those given above for all-atom simulations of CPTP bound to a membrane. 300 ns production runs were then performed for each bilayer. The final configuration from the simulation of the membrane composed of both POPC and 16:0-C1P was used to initialize one enhanced sampling simulation. Another initial configuration was constructed by (1) inserting a 16:0-C1P lipid randomly into the solution above the POPC membrane such that its COM was at least 3.2 nm above the COM of the membrane, (2) energy minimizing the system, and (3) equilibrating it using the two-step protocol used for the membrane only system with the addition of harmonic restraints with a force constant of 500 kJ/mol/nm^2^ on the *z* coordinates of all heavy atoms of 16:0-C1P. Next, to generate configurations to initialize each umbrella sampling window, 20 ns steered MD simulations were performed using a harmonic bias on *r*_LxS_ with a spring constant of 75 kJ/mol. During one steered simulation, a tagged 16:0-C1P was pulled from *r*_LxS_ = 14 to *r*_LxS_ = −5 to extract it from the membrane. In the other, the 16:0-C1P was pulled from *r*_LxS_ = −5 to *r*_LxS_ = 14 to insert it into the membrane. A total of 66 umbrella sampling windows were simulated with equally spaced harmonic bias centers ranging from *r*_LxS_ = −5 to *r*_LxS_ = 15 and a spring constant of 40 kJ/mol. All windows were run for 52 ns (amounting to a cumulative simulation time of 3.4 *μ*s per system), and the first 20 ns was discarded to account for equilibration. The weighted histogram analysis method (WHAM) [95] was used to combine data from all windows and obtain a free energy profile. Free energy profiles calculated for the two independent systems are in agreement, indicating convergence (S11 Fig). Data from all windows of both systems was then combined to obtain the free energy profile for passive lipid transport shown in Fig 7B, and error bars were calculated as the standard error estimated from the two independent systems.

## Supporting information

**S1 Fig. In solution, one side of CPTP’s sandwich-like structure adjusts to the presence of C1P.** Root-mean-square deviation (RMSD) of each residue’s C*α* atom of CPTP in its apo form relative to its C1P-bound form during solution-phase all-atom simulations. After alignment of the structures to minimize the RMSD of C*α* atoms of helix *α*6, the RMSD between each C*α* in the C1P-bound and apo forms is calculated as 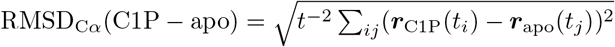, where *i* and *j* index all *t* frames in each trajectory and *r*_c1P_(*t*) and *r*_apo_(*t*) are the C*α*’s positions in the C1P-bound and apo forms, respectively. CPTP’s secondary structure is schematically illustrated above with helices represented as rectangles and unstructured loop regions as lines. The gray region highlights residues that comprise one side of CPTP’s sandwich-like structure. The dashed lines indicated the average RMSD_c*α*_(C1P – apo) for helices *α*N and *α*1 − 3 of 4.6 Å (white region) and helices *α*4, *α*5, *α*7, and *α*8 of 3.6 Å (gray region).

**S2 Fig. Membrane binding is observed in unbiased coarse-grained simulations.** (A) Minimum distance between CPTP and the membrane during coarse-grained simulations of membrane binding. (B) Configurations observed at the end of each simulation. Four independent simulations were performed for each form of CPTP. Within 2 *μ*s, CPTP stably bound the membrane in all but one simulation of the apo form (indicated with an asterisk).

**S3 Fig. Coarse-grained representations of CPTP reproduce its internal dynamics observed in all-atom simulations.** Root-mean-square fluctuation (RMSF) of C*α* atoms (backbone beads) of CPTP in its apo and C1P-bound forms during solution-phase all-atom (coarse-grained) simulations.

**S4 Fig. Coarse-grained representations of CPTP match its structure in all-atom simulations.** Root-mean-square deviation (RMSD) of C*α* atoms (backbone beads) between CPTP in its apo and C1P-bound forms during solution-phase all-atom (coarse-grained) simulations and the cystral structure of CPTP in PDB 4K85. In the top row, the RMSD for residues 8 – 214 is plotted. All other rows show the RMSD for individual helices.

**S5 Fig. CPTP alters the membrane’s physical properties to facilitate extraction and insertion of C1P into the membrane.** CPTP impacts (A and B) the average orientational order parameter of lipids’ acyl chains, *S*_CC_ [84], (C and D) membrane thickness, and (E and F) area per lipid. (A, C, and E) Average of each property as a function of a lipid’s displacement in the *xy* plane from CPTP. Color scales are set relative to the average value for a POPC membrane without CPTP present (white). Structures of the apo and C1P-bound forms of CPTP bound to the membrane are outlined. (B, D, and F) Box-and-whisker plots of each property for lipids within a specified radial distance from CPTP’s center-of-mass in the xy plane and for lipids in a POPC membrane without CPTP present. The box extends from the 1st to 3rd quartile with the median indicated by the black line, and the whiskers extend from the minimum to maximum value. Asterisks indicate the average differs significantly from that of a POPC membrane: ** *p* < 0.001; * *p* < 0.01 (Welch’s t-test).

**S6 Fig. Conformational differences between the apo and C1P-bound forms of CPTP result in differing effects on the membrane’s physical properties.** In both the apo and C1P-bound forms, residues on helices *α*2 and *α*6 promote lipid reorientation. Residues on helix α5 of the C1P-bound form promote further increases in lipid reorientation and disorder. Residues on helices *α*1, *α*2, and *α*6, especially in the the apo form of CPTP, promote decreases in local membrane hydrophobicity and increases in the area per lipid. Average change in (A) cos(*θ_z_*) (Fig 5A), (B) *S*_CC_ [84], (C) *n*_CC_ (Fig 6A), and (D) area per lipid, *A*_lip_, relative to the average of a POPC membrane without CPTP present for lipids within 5 Å of each residue of CPTP. CPTP’s secondary structure is schematically illustrated above with helices represented as rectangles and unstructed loop regions as lines.

**S7 Fig. CPTP forms hydrophobic contacts with lipids to disrupt their hydrophobic membrane environments.** 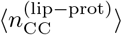 mapped onto the structures of the (A-E) apo and (F-J) C1P-bound forms. Example configurations of lipids with (1) a reduction in *n*_CC_ of ≥ 300 contacts relative to 〈*n*_CC_〉_mem_ of a POPC membrane without CPTP bound and (2) ≥ 90% of those contacts replaced with hydrophobic contacts with CPTP are shown. The black line indicates the average position of phosphate groups of membrane lipids.

**S8 Fig. Multi-walker well-tempered metadynamics simulation converges after 500 ns.** (A and B) Walkers initiated from the apo (cyan points outlined in gray) and C1P-bound forms (magenta points outlined in gray) broadly and diffusively sample the space of *r*_LxS_ and *Q* and are well-mixed by the end of the simulation. Values of *r*_LxS_ and *Q* at the end of the simulation are indicated with points outlined in black for each walker (cyan for those initialized from the apo form and magenta for those initialized from the C1P-bound form). (A) Values of *r*_LxS_ and *Q* sampled during the simulation. Each point is colored by time with configurations sampled at early times shown in dark blue and at later times in dark red. (B) Trajectories of *r*_LxS_ and *Q* for each walker. Walkers initiated from the apo form are plotted in cool colors, and walkers initiated from the C1P-bound form are plotted in warm colors. (C) Scaled time computed from the time-dependent bias is plotted versus simulation time on a log-log scale. The dashed gray line is the linear fit after 500 ns. The slope of this line is 29, which is consistent with the bias factor of 30 used in the simulation and suggests that a quasi-steady state has been reached. (D and E) Agreement between free energy profiles calculated at simulation times ranging from 760 ns (cumulatively 15.2 *μ*s) to 1140 ns (cumulatively 22.8 *μ*s) indicates convergence. All free energy profiles were calculated by reweighting frames after 500 ns (cumulatively 10 *μ*s) up to the specified time using the estimator of Tiwary and Parrinello [91].

**S9 Fig. Error in the free energy surface Δ*F*(*r*_LxS_,*Q*)**. (A) The free energy surface Δ*F*(*r*_LxS_, *Q*) shown in Fig 7A is reproduced. (B) Standard error of Δ*F*(*r*_LxS_, *Q*) computed with block averaging.

**S10 Fig. Free energy profile as a function of *Q***. Δ*F*(*Q*) is obtained by marginalizing Δ*F*(*r*_LxS_,*Q*) over *r*_LxS_. Error bars indicate the standard error computed with block averaging.

**S11 Fig. Agreement between free energy profiles of passive C1P transport calculated for two independent systems indicates convergence of umbrella sampling simulations.** Free energy profiles along *r*_LxS_ calculated for two independent systems. For each system’s free energy profile, error bars were calculated as the standard error of Δ*F* estimated from four independent 8 ns blocks.

**S12 Fig. Many different intermediate configurations are sampled during CPTP extraction (or insertion) of C1P from (or into) a membrane.** View looking into CPTP’s hydrophobic cavity of example configurations from different regions of the free energy surface outlined in Fig 7A. C1P is rendered as van der Waals spheres and colored orange. The apo form of CPTP in configuration 1 is colored dark cyan, and the C1P-bound form in configuration 12 is colored dark magenta. The black line indicates the average position of phosphate groups of membrane lipids.

**S1 Text. Supporting Notes.** Discussion of (1) conformational differences between apo and C1P-bound forms of CPTP both in solution-phase and membrane-bound simulations analyzed with principal component analysis and (2) choice of C1P–CPTP contacts used to define *Q*.

## Author contributions

Conceptualization: J.R.R.; Data curation: J.R.R.; Formal analysis: J.R.R.; Investigation: J.R.R.; Methodology: J.R.R., P.L.G.; Resources: J.R.R., P.L.G.; Software: J.R.R.; Supervision: P.L.G.; Visualization: J.R.R.; Writing – Original draft preparation: J.R.R.; Writing – Review & editing: J.R.R., P.L.G

## Data availability

The data that support the findings of this study are available in the manuscript and its supporting information files. All simulation inputs, equilibrium simulation trajectories, and analysis results are available on Zenodo: https://doi.org/10.5281/zenodo.7067163. Analysis code is available on GitHub: https://github.com/jrurogers/cptp-analysis.

## Acknowledgements

We thank David Limmer and Layne Frechette for valuable discussions and useful comments on the manuscript. This work is dedicated to the memory of late Prof. Phill Geissler, a cherished teacher, mentor, and colleague—an inspiration.

## References

1. Hannun YA, Obeid LM. Sphingolipids and their metabolism in physiology and disease. Nat Rev Mol Cell Biol. 2018;19(3):175–191.

2. Olson DK, Frohlich F, Farese RV, Walther TC. Taming the sphinx: Mechanisms of cellular sphingolipid homeostasis. Biochim Biophys Acta. 2016;1861(8):784–792.

3. Yamaji T, Hanada K. Sphingolipid metabolism and interorganellar transport: Localization of sphingolipid enzymes and lipid transfer proteins. Traffic. 2015;16(2):101–122.

4. Breslow DK. Sphingolipid homeostasis in the endoplasmic reticulum and beyond. Cold Spring Harb Perspect Biol. 2013;5(4):a013326–a013326.

5. Maceyka M, Spiegel S. Sphingolipid metabolites in inflammatory disease. Nature. 2014;510(7503):58–67.

6. Ogretmen B. Sphingolipid metabolism in cancer signalling and therapy. Nat Rev Cancer. 2018;18:33–50.

7. Samaha D, Hamdo HH, Wilde M, Prause K, Arenz C. Sphingolipid-transporting proteins as cancer therapeutic targets. Int J Mol Sci. 2019;20(14):3554.

8. Körner C, Fröhlich F. Compartmentation and functions of sphingolipids. Curr Opin Cell Biol. 2022;74:104–111.

9. Backman APE, Mattjus P. Who moves the sphinx? An overview of intracellular sphingolipid transport. Biochim Biophys Acta Mol Cell Biol Lipids. 2021;1866(11):159021.

10. Reinisch KM, Prinz WA. Mechanisms of nonvesicular lipid transport. J Cell Biol. 2021;220(3):e202012058.

11. Egea PF. Mechanisms of non-vesicular exchange of lipids at membrane contact sites: Of shuttles, tunnels and, funnels. Front Cell Dev Biol. 2021;9:784367.

12. Jackson CL, Walch L, Verbavatz JM. Lipids and their trafficking: An integral part of cellular organization. Dev Cell. 2016;39(2):139–153.

13. Lev S. Non-vesicular lipid transport by lipid-transfer proteins and beyond. Nat Rev Mol Cell Biol. 2010;11:739–750.

14. Wong LH, Gatta AT, Levine TP. Lipid transfer proteins: The lipid commute via shuttles, bridges and tubes. Nat Rev Mol Cell Biol. 2019;20(2):85–101.

15. Arana L, Gangoiti P, Ouro A, Trueba M, Gómez-Muñoz A. Ceramide and ceramide 1-phosphate in health and disease. Lipids Health Dis. 2010;9:15.

16. Simanshu DK, Kamlekar RK, Wijesinghe DS, Zou X, Zhai X, Mishra SK, et al. Non-vesicular trafficking by a ceramide-1-phosphate transfer protein regulates eicosanoids. Nature. 2013;500(7463):463–467.

17. Mishra SK, Gao YG, Deng YB, Chalfant CE, Hinchcliffe EH, Brown RE. CPTP: A sphingolipid transfer protein that regulates autophagy and inflammasome activation. Autophagy. 2018;14(5):862–879.

18. Mishra SK, Gao YG, Zou X, Stephenson DJ, Malinina L, Hinchcliffe EH, et al. Emerging roles for human glycolipid transfer protein superfamily members in the regulation of autophagy, inflammation, and cell death. Prog Lipid Res. 2020;78:101031.

19. Zhai XH, Gao YG, Mishra SK, Simanshu DK, Boldyrev IA, Benson LM, et al. Phosphatidylserine stimulates ceramide-1-phosphate (C1P) intermembrane transfer by C1P transfer proteins. J Biol Chem. 2017;292(6):2531–2541.

20. Gao YG, Zhai X, Boldyrev IA, Molotkovsky JG, Patel DJ, Malinina L, et al. Ceramide-1-phosphate transfer protein (CPTP) regulation by phosphoinositides. J Biol Chem. 2021;296:100600.

21. McLean LR, Phillips MC. Kinetics of phosphatidylcholine and lysophosphatidylcholine exchange between unilamellar vesicles. Biochemistry. 1984;23(20):4624–4630.

22. Nichols JW. Thermodynamics and kinetics of phospholipid monomer vesicle interaction. Biochemistry. 1985;24(23):6390–6398.

23. Pownall HJ, Hickson DL, Smith LC. Transport of biological lipophiles: Effect of lipophile structure. J Am Chem Soc. 1983;105(8):2440–2445.

24. Xia Y, Li M, Charubin K, Liu Y, Heberle FA, Katsaras J, et al. Effects of nanoparticle morphology and acyl chain length on spontaneous lipid transfer rates. Langmuir. 2015;31(47):12920–12928.

25. Rogers JR, Geissler PL. Breakage of hydrophobic contacts limits the rate of passive lipid exchange between membranes. J Phys Chem B. 2020;124(28):5884–5898.

26. Rogers JR, Espinoza Garcia G, Geissler PL. Membrane hydrophobicity determines the activation free energy of passive lipid transport. Biophys J. 2021;120(17):3718–3731.

27. Gao YG, McDonald J, Malinina L, Patel DJ, Brown RE. Ceramide-1-phosphate transfer protein promotes sphingolipid reorientation needed for binding during membrane interaction. J Lipid Res. 2022;63(1):100151.

28. Malinina L, Patel DJ, Brown RE. How α-helical motifs form functionally diverse lipid-binding compartments. Annu Rev Biochem. 2017;86(1):609–636.

29. Malinina L, Simanshu DK, Zhai X, Samygina VR, Kamlekar R, Kenoth R, et al. Sphingolipid transfer proteins defined by the GLTP-fold. Q Rev Biophys. 2015;48(3):281–322.

30. Brown RE, Mattjus P. Glycolipid transfer proteins. Biochim Biophys Acta Mol Cell Biol Lipids. 2007;1771(6):746–760.

31. Schmidtke P, Bidon-Chanal A, Luque FJ, Barril X. MDpocket: Open-source cavity detection and characterization on molecular dynamics trajectories. Bioinformatics. 2011;27(23):3276–3285.

32. Bolhuis PG, Chandler D, Dellago C, Geissler PL. Transition path sampling: Throwing ropes over rough mountain passes, in the dark. Annu Rev Phys Chem. 2002;53:291–318.

33. Du R, Pande VS, Grosberg AY, Tanaka T, Shakhnovich ES. On the transition coordinate for protein folding. J Chem Phys. 1998;108(1):334–350.

34. Zuckerman DM. Statistical physics of biomolecules: An introduction. Boca Raton, FL: CRC Press; 2010.

35. Best RB, Hummer G, Eaton WA. Native contacts determine protein folding mechanisms in atomistic simulations. Proc Natl Acad Sci U S A. 2013;110(44):17874–17879.

36. Raiteri P, Laio A, Gervasio FL, Micheletti C, Parrinello M. Efficient reconstruction of complex free energy landscapes by multiple walkers metadynamics. J Phys Chem B. 2006;110(8):3533–3539.

37. Barducci A, Bussi G, Parrinello M. Well-tempered metadynamics: A smoothly converging and tunable free-energy method. Phys Rev Lett. 2008;100(2).

38. Pabis A, Rawle RJ, Kasson PM. Influenza hemagglutinin drives viral entry via two sequential intramembrane mechanisms. Proc Natl Acad Sci U S A. 2020;117(13):7200–7207.

39. Holthuis JCM, Menon AK. Lipid landscapes and pipelines in membrane homeostasis. Nature. 2014;510:48–57.

40. Leonard AN, Simmonett AC, Pickard FC IV, Huang J, Venable RM, Klauda JB, et al. Comparison of additive and polarizable models with explicit treatment of long-range Lennard-Jones interactions using alkane simulations. J Chem Theory Comput. 2018;14(2):948–958.

41. Kramer A, Pickard FC, Huang J, Venable RM, Simmonett AC, Reith D, et al. Interactions of water and alkanes: Modifying additive force fields to account for polarization effects. J Chem Theory Comput. 2019;15(6):3854–3867.

42. Sega M, Dellago C. Long-range dispersion effects on the water/vapor interface simulated using the most common models. J Phys Chem B. 2017;121(15):3798–3803.

43. Simanshu D, Zhai X, Munch D, Hofius D, Markham J, Bielawski J, et al. Arabidopsis accelerated cell death 11, ACD11, is a ceramide-1-phosphate transfer protein and intermediary regulator of phytoceramide levels. Cell Rep. 2014;6(2):388–399.

44. Harayama T, Riezman H. Understanding the diversity of membrane lipid composition. Nat Rev Mol Cell Biol. 2018;19(5):281–296.

45. Pogozheva ID, Armstrong GA, Kong L, Hartnagel TJ, Carpino CA, Gee SE, et al. Comparative molecular dynamics simulation studies of realistic eukaryotic, prokaryotic, and archaeal membranes. J Chem Inf Model. 2022;62(4):1036–1051.

46. Ingólfsson HI, Melo MN, van Eerden FJ, Arnarez C, Lopez CA, Wassenaar TA, et al. Lipid organization of the plasma membrane. J Am Chem Soc. 2014;136(41):14554–14559.

47. Wong LH, Copic A, Levine TP. Advances on the transfer of lipids by lipid transfer proteins. Trends Biochem Sci. 2017;42(7):516–530.

48. Miliara X, Tatsuta T, Berry JL, Rouse SL, Solak K, Chorev DS, et al. Structural determinants of lipid specificity within Ups/PRELI lipid transfer proteins. Nat Commun. 2019;10:1130.

49. Watanabe Y, Tamura Y, Kawano S, Endo T. Structural and mechanistic insights into phospholipid transfer by Ups1–Mdm35 in mitochondria. Nat Commun. 2015;6:7922.

50. Grabon A, Orlowski A, Tripathi A, Vuorio J, Javanainen M, Rog T, et al. Dynamics and energetics of the mammalian phosphatidylinositol transfer protein phospholipid exchange cycle. J Biol Chem. 2017;292(35):14438–14455.

51. Iaea DB, Dikiy I, Kiburu I, Eliezer D, Maxfield FR. STARD4 membrane interactions and sterol binding. Biochemistry. 2015;54(30):4623–4636.

52. Kudo N, Kumagai K, Tomishige N, Yamaji T, Wakatsuki S, Nishijima M, et al. Structural basis for specific lipid recognition by CERT responsible for nonvesicular trafficking of ceramide. Proc Natl Acad Sci U S A. 2008;105(2):488–493.

53. Sugiura T, Takahashi C, Chuma Y, Fukuda M, Yamada M, Yoshida U, et al. Biophysical parameters of the Sec14 phospholipid exchange cycle. Biophys J. 2019;116(1):92–103.

54. Dong J, Du X, Wang H, Wang J, Lu C, Chen X, et al. Allosteric enhancement of ORP1-mediated cholesterol transport by PI(4,5)P2/PI(3,4)P2. Nat Commun. 2019;10:829.

55. Lipp NF, Gautier R, Magdeleine M, Renard M, Albanèse V, Čopič A, et al. An electrostatic switching mechanism to control the lipid transfer activity of Osh6p. Nat Commun. 2019;10:3926.

56. Shadan S, Holic R, Carvou N, Ee P, Li M, Murray-Rust J, et al. Dynamics of lipid transfer by phosphatidylinositol transfer proteins in cells. Traffic. 2008;9(10):1743–1756.

57. Zhang X, Xie H, Iaea D, Khelashvili G, Weinstein H, Maxfield FR. Phosphatidylinositol phosphates modulate interactions between the StarD4 sterol trafficking protein and lipid membranes. J Biol Chem. 2022;298(7):102058.

58. Abraham MJ, Murtola T, Schulz R, Páll S, Smith JC, Hess B, et al. GROMACS: High performance molecular simulations through multi-level parallelism from laptops to supercomputers. SoftwareX. 2015;1-2:19–25.

59. Klauda JB, Venable RM, Freites JA, O’Connor JW, Tobias DJ, Mondragon-Ramirez C, et al. Update of the CHARMM all-atom additive force field for lipids: Validation on six lipid types. J Phys Chem B. 2010;114(23):7830–7843.

60. MacKerell AD, Bashford D, Bellott M, Dunbrack RL, Evanseck JD, Field MJ, et al. All-atom empirical potential for molecular modeling and dynamics studies of proteins. J Phys Chem B. 1998;102(18):3586–3616.

61. Venable RM, Sodt AJ, Rogaski B, Rui H, Hatcher E, MacKerell AD, et al. CHARMM all-atom additive force field for sphingomyelin: Elucidation of hydrogen bonding and of positive curvature. Biophys J. 2014;107(1):134–145.

62. Khan HM, Souza PCT, Thallmair S, Barnoud J, de Vries AH, Marrink SJ, et al. Capturing choline-aromatics cation-pi interactions in the MARTINI force field. J Chem Theory Comput. 2020;16(4):2550–2560.

63. Yesylevskyy SO, Schafer LV, Sengupta D, Marrink SJ. Polarizable water model for the coarse-grained MARTINI force field. PLoS Comput Biol. 2010;6(6):e1000810.

64. Periole X, Cavalli M, Marrink SJ, Ceruso MA. Combining an elastic network with a coarse-grained molecular force field: Structure, dynamics, and intermolecular recognition. J Chem Theory Comput. 2009;5(9):2531–2543.

65. Marrink SJ, Risselada HJ, Yefimov S, Tieleman DP, de Vries AH. The MARTINI force field: Coarse grained model for biomolecular simulations. J Phys Chem B. 2007;111(27):7812–7824.

66. Webb B, Sali A. Comparative protein structure modeling using MODELLER. Curr Protoc Bioinform. 2016;54:5.6.1–5.6.37.

67. Fiser A, Do RKG, Sali A. Modeling of loops in protein structures. Protein Sci. 2000;9(9):1753–1773.

68. Berendsen HJC, Postma JPM, van Gunsteren WF, DiNola A, Haak JR. Molecular dynamics with coupling to an external bath. J Chem Phys. 1984;81(8):3684–3690.

69. Parrinello M, Rahman A. Polymorphic transitions in single-crystals - a new molecular-dynamics method. J Appl Phys. 1981;52(12):7182–7190.

70. Wassenaar TA, Ingólfsson HI, Böckmann RA, Tieleman DP, Marrink SJ. Computational lipidomics with insane: A versatile tool for generating custom membranes for molecular simulations. J Chem Theory Comput. 2015;11(5):2144–2155.

71. de Jong DH, Singh G, Bennett WFD, Arnarez C, Wassenaar TA, Schafer LV, et al. Improved parameters for the martini coarse-grained protein force field. J Chem Theory Comput. 2013;9:687–697.

72. Wassenaar TA, Pluhackova K, Bockmann RA, Marrink SJ, Tieleman DP. Going backward: A flexible geometric approach to reverse transformation from coarse grained to atomistic models. J Chem Theory Comput. 2014;10(2):676–690.

73. Nosé S. A molecular-dynamics method for simulations in the canonical ensemble. Mol Phys. 1984;52(2):255–268.

74. Hoover WG. Canonical dynamics - equilibrium phase-space distributions. Phys Rev A. 1985;31(3):1695–1697.

75. Hockney WR. The potential calculation and some applications. Methods Comput Phys. 1970;9:136–211.

76. Hess B, Bekker H, Berendsen HJ, Fraaije JG. Lincs: A linear constraint solver for molecular simulations. J Comput Chem. 1997;18(12):1463–1472.

77. Essmann U, Perera L, Berkowitz ML, Darden T, Lee H, Pedersen LG. A smooth particle mesh ewald method. J Chem Phys. 1995;103(19):8577–8593.

78. Páll S, Hess B. A flexible algorithm for calculating pair interactions on SIMD architectures. Comput Phys Commun. 2013;184(12):2641–2650.

79. Lingenheil M, Denschlag R, Reichold R, Tavan P. The “hot-solvent/cold-solute” problem revisited. J Chem Theory Comput. 2008;4(8):1293–306.

80. Bussi G, Donadio D, Parrinello M. Canonical sampling through velocity rescaling. J Chem Phys. 2007;126(1):014101.

81. Michaud-Agrawal N, Denning EJ, Woolf TB, Beckstein O. MDAnalysis: A toolkit for the analysis of molecular dynamics simulations. J Comput Chem. 2011;32(10):2319–2327.

82. Harris CR, Millman KJ, van der Walt SJ, Gommers R, Virtanen P, Cournapeau D, et al. Array programming with numpy. Nature. 2020;585(7825):357–362.

83. Buchoux S. FATSLiM: A fast and robust software to analyze MD simulations of membranes. Bioinformatics. 2017;33(1):133–134.

84. Seo S, Murata M, Shinoda W. Pivotal role of interdigitation in interleaflet interactions: Implications from molecular dynamics simulations. J Phys Chem Lett. 2020;11(13):5171–5176.

85. Smith P, Lorenz CD. LiPyphilic: A python toolkit for the analysis of lipid membrane simulations. J Chem Theory Comput. 2021;17(9):5907–5919.

86. Flyvbjerg H, Petersen HG. Error estimates on averages of correlated data. J Chem Phys. 1989;91:461–466.

87. Welch BL. The generalization of ‘student’s’ problem when several different population varlances are involved. Biometrika. 1947;34(1-2):28–35.

88. Bochicchio D, Panizon E, Ferrando R, Monticelli L, Rossi G. Calculating the free energy of transfer of small solutes into a model lipid membrane: Comparison between metadynamics and umbrella sampling. J Chem Phys. 2015;143(14):144108.

89. Prajapati JD, Fernández Solano CJ, Winterhalter M, Kleinekathöfer U. Characterization of ciprofloxacin permeation pathways across the porin OmpC using metadynamics and a string method. J Chem Theory Comput. 2017;13(9):4553–4566.

90. Tribello GA, Bonomi M, Branduardi D, Camilloni C, Bussi G. PLUMED 2: New feathers for an old bird. Comput Phys Commun. 2014;185(2):604–613.

91. Tiwary P, Parrinello M. A time-independent free energy estimator for metadynamics. J Phys Chem B. 2015;119(3):736–742.

92. Torrie GM, Valleau JP. Non-physical sampling distributions in monte-carlo free-energy estimation - umbrella sampling. J Comput Phys. 1977;23(2):187–199.

93. Jo S, Kim T, Iyer VG, Im W. CHARMM-GUI: A web-based graphical user interface for CHARMM. J Comput Chem. 2008;29(11):1859–1865.

94. Wu EL, Cheng X, Jo S, Rui H, Song KC, Dávila-Contreras EM, et al. CHARMM-GUI membrane builder toward realistic biological membrane simulations. J Comput Chem. 2014;35(27):1997–2004.

95. Kumar S, Bouzida D, Swendsen RH, Kollman PA, Rosenberg JM. The weighted histogram analysis method for free-energy calculations on biomolecules. 1. The method. J Comput Chem. 1992;13(8):1011–1021.

