## Supporting information for "Ceramide-1-phosphate transfer protein enhances lipid transport by disrupting hydrophobic lipid–membrane contacts"

### SUPPORTING FIGURES

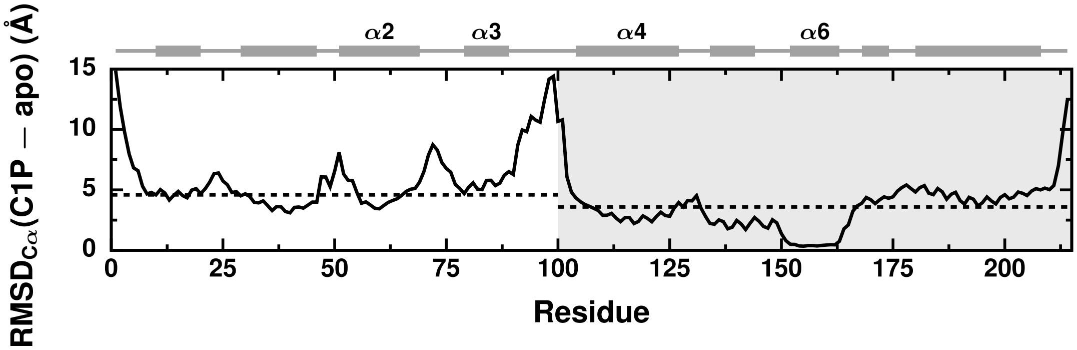


**Fig S1. In solution, one side of CPTP's sandwich-like structure adjusts to the presence of C1P.** Root-mean-square deviation (RMSD) of each residue's C$\alpha$ atom of CPTP in its apo form relative to its C1P-bound form during solution-phase all-atom simulations. After alignment of the structures to minimize the RMSD of C$\alpha$ atoms of helix $\alpha 6$, the RMSD between each C$\alpha$ in the C1P-bound and apo forms is calculated as $\mathrm{RMSD}_{\mathrm{C}\alpha}(\mathrm{C1P}-\mathrm{apo}) = \sqrt{t^{-2}\sum_{ij}(\boldsymbol{r}_{\mathrm{C1P}}(t_i)-\boldsymbol{r}_{\mathrm{apo}}(t_j))^2}$, where $i$ and $j$ index all $t$ frames in each trajectory and $\boldsymbol{r}_{\mathrm{C1P}}(t)$ and $\boldsymbol{r}_{\mathrm{apo}}(t)$ are the C$\alpha$'s positions in the C1P-bound and apo forms, respectively. CPTP's secondary structure is schematically illustrated above with helices represented as rectangles and unstructured loop regions as lines. The gray region highlights residues that comprise one side of CPTP's sandwich-like structure. The dashed lines indicated the average $\mathrm{RMSD}_{\mathrm{C}\alpha}(\mathrm{C1P}-\mathrm{apo})$ for helices $\alpha$N and $\alpha 1-3$ of 4.6 Å (white region) and helices $\alpha 4, \alpha 5, \alpha 7$, and $\alpha 8$ of 3.6 Å (gray region).

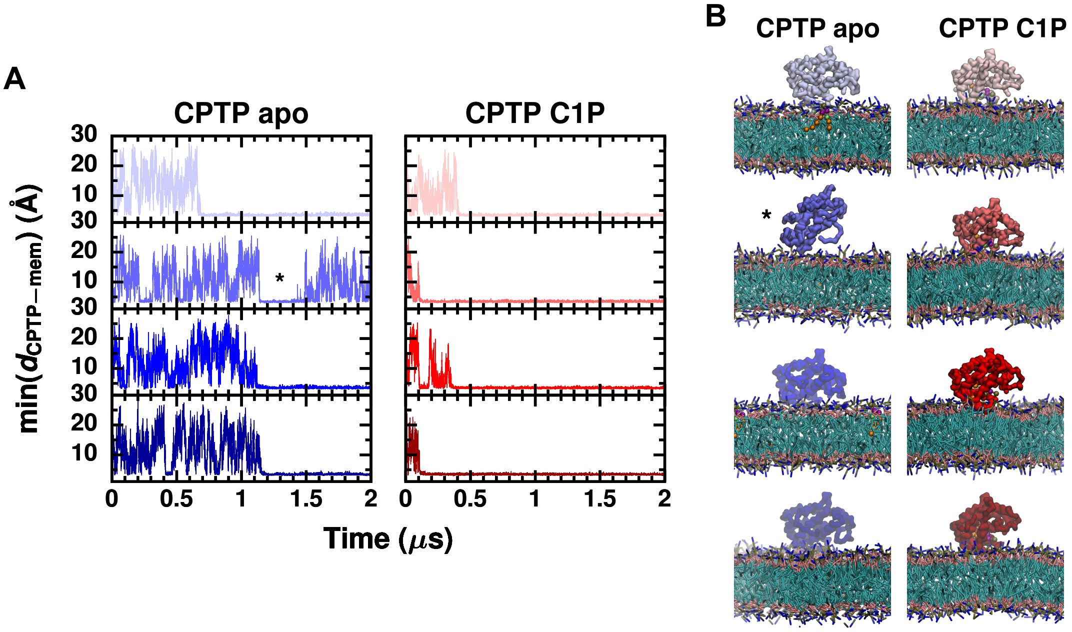


**Fig S2. Membrane binding is observed in unbiased coarse-grained simulations.** (A) Minimum distance between CPTP and the membrane during coarse-grained simulations of membrane binding. (B) Configurations observed at the end of each simulation. Four independent simulations were performed for each form of CPTP. Within 2 $\mu$s, CPTP stably bound the membrane in all but one simulation of the apo form (indicated with an asterisk).

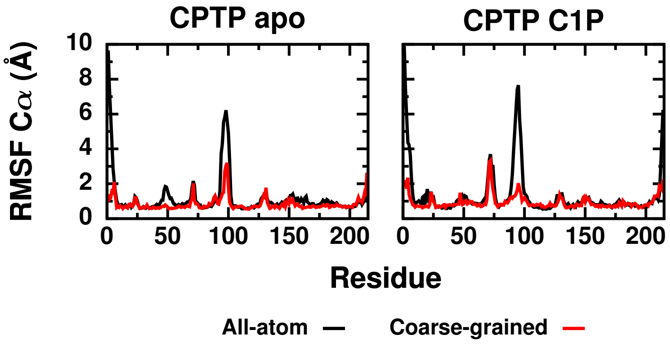


**Fig S3. Coarse-grained representations of CPTP reproduce its internal dynamics observed in all-atom simulations.** Root-mean-square fluctuation (RMSF) of C$\alpha$ atoms (backbone beads) of CPTP in its apo and C1P-bound forms during solution-phase all-atom (coarse-grained) simulations.

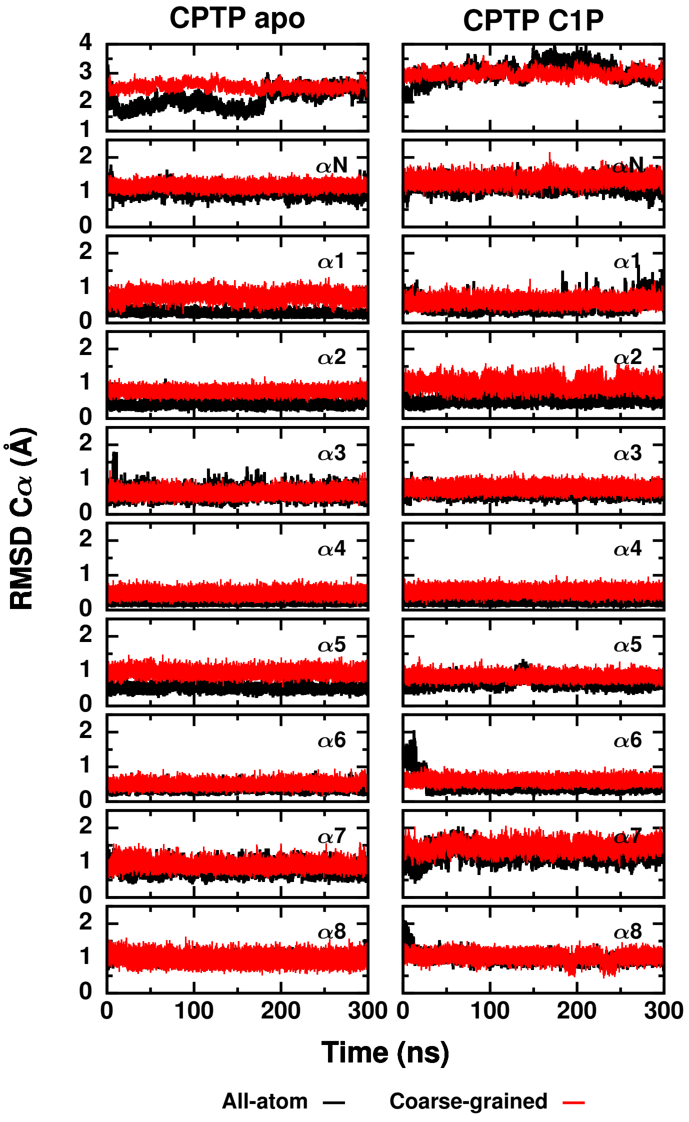


**Fig S4. Coarse-grained representations of CPTP match its structure in all-atom simulations.** Root-mean-square deviation (RMSD) of C$\alpha$ atoms (backbone beads) between CPTP in its apo and C1P-bound forms during solution-phase all-atom (coarse-grained) simulations and the cystral structure of CPTP in PDB 4K85. In the top row, the RMSD for residues $8-214$ is plotted. All other rows show the RMSD for individual helices.

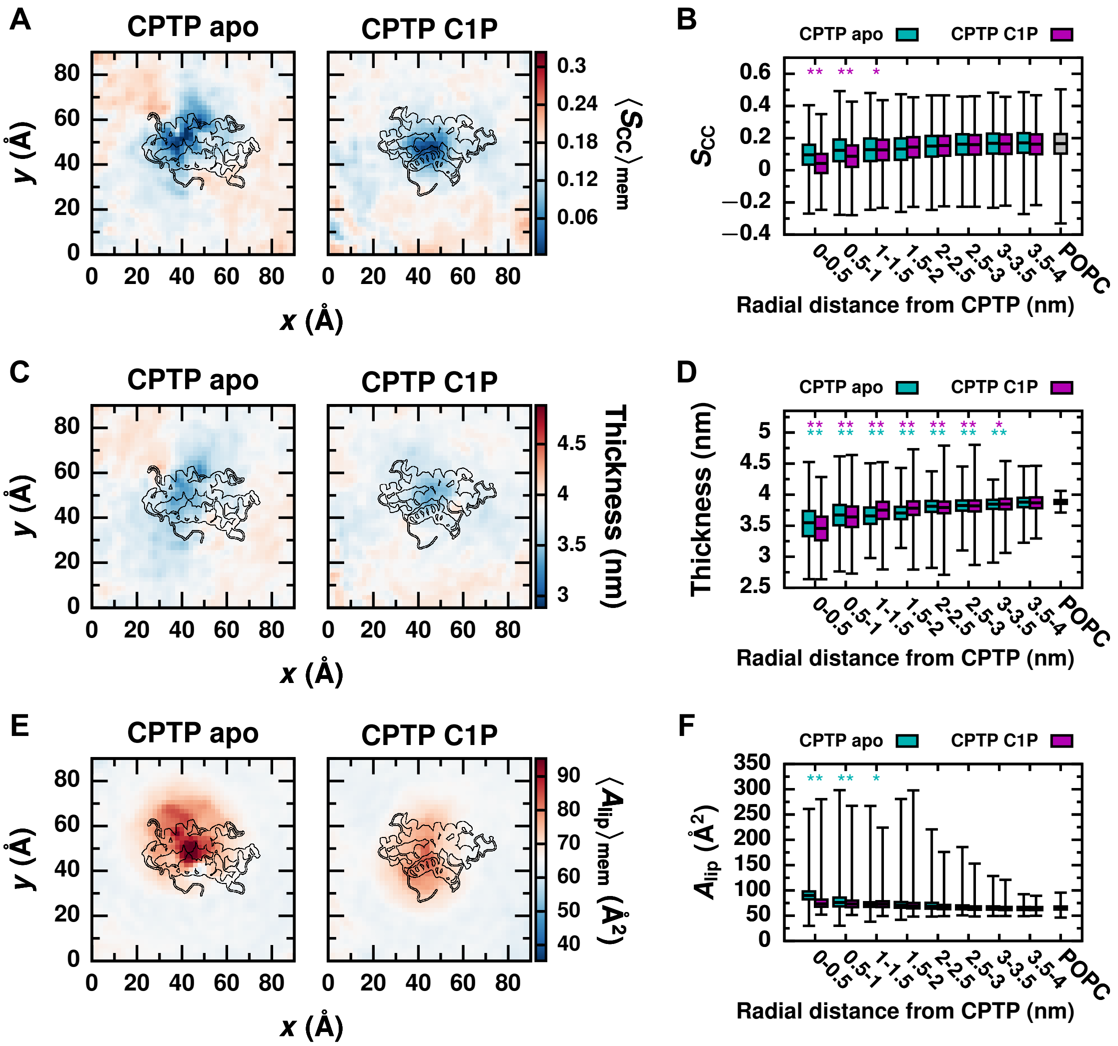


**Fig S5. CPTP alters the membrane's physical properties to facilitate extraction and insertion of C1P into the membrane.** CPTP impacts (A and B) the average orientational order parameter of lipids' acyl chains, $S_{\mathrm{CC}}$ [84], (C and D) membrane thickness, and (E and F) area per lipid. (A, C, and E) Average of each property as a function of a lipid's displacement in the $xy$ plane from CPTP. Color scales are set relative to the average value for a POPC membrane without CPTP present (white). Structures of the apo and C1P-bound forms of CPTP bound to the membrane are outlined. (B, D, and F) Box-and-whisker plots of each property for lipids within a specified radial distance from CPTP's center-of-mass in the $xy$ plane and for lipids in a POPC membrane without CPTP present. The box extends from the 1st to 3rd quartile with the median indicated by the black line, and the whiskers extend from the minimum to maximum value. Asterisks indicate the average differs significantly from that of a POPC membrane: ** $p < 0.001$; * $p < 0.01$ (Welch's t-test).

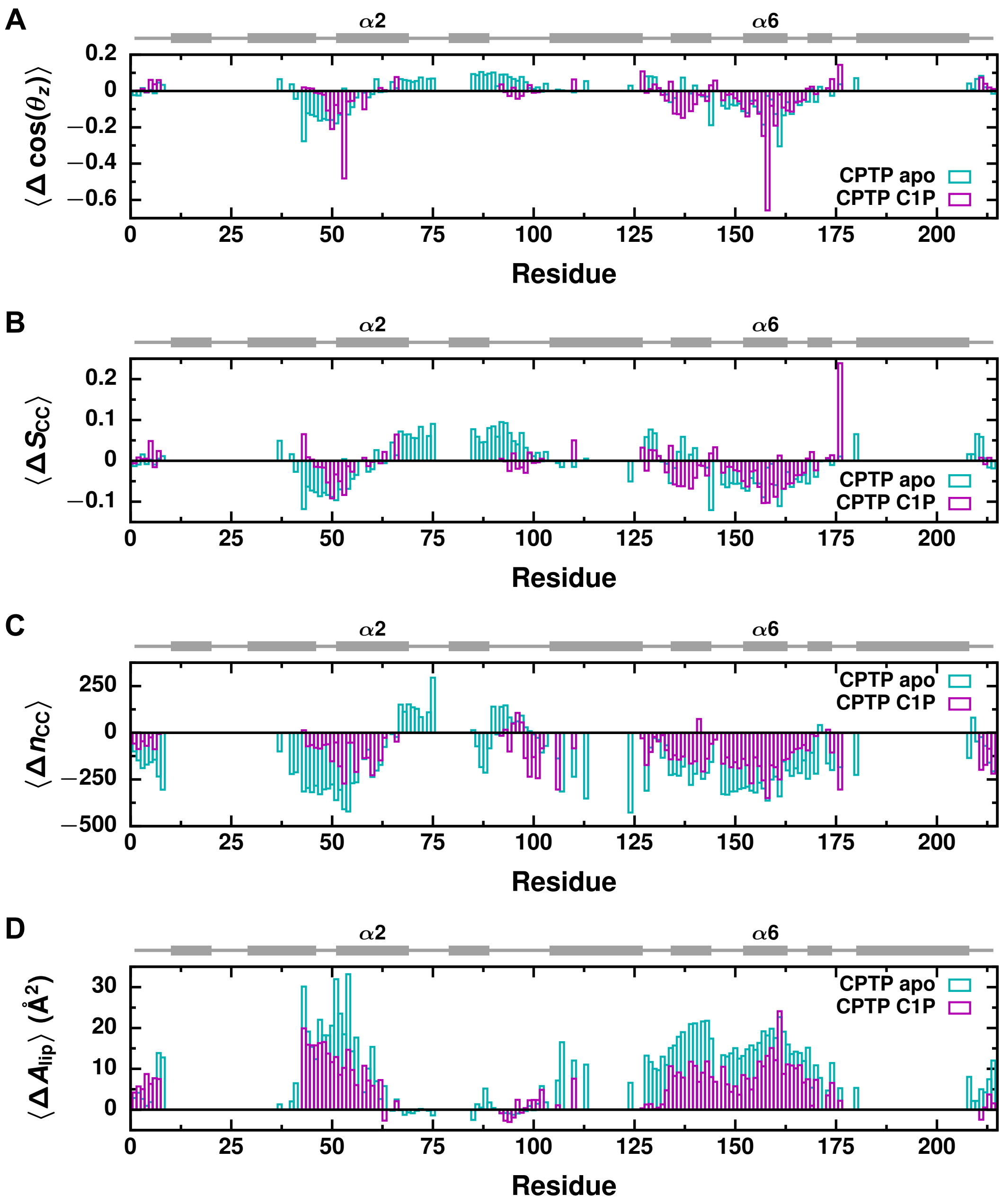


**Fig S6. Conformational differences between the apo and C1P-bound forms of CPTP result in differing effects on the membrane's physical properties.** In both the apo and C1P-bound forms, residues on helices $\alpha 2$ and $\alpha 6$ promote lipid reorientation. Residues on helix $\alpha 5$ of the C1P-bound form promote further increases in lipid reorientation and disorder. Residues on helices $\alpha 1$, $\alpha 2$, and $\alpha 6$, especially in the the apo form of CPTP, promote decreases in local membrane hydrophobicity and increases in the area per lipid. Average change in (A) $\cos(\theta_z)$ (Fig 5A), (B) $S_{\mathrm{CC}}$ [84], (C) $n_{\mathrm{CC}}$ (Fig 6A), and (D) area per lipid, $A_{\mathrm{lip}}$, relative to the average of a POPC membrane without CPTP present for lipids within 5Å of each residue of CPTP. CPTP's secondary structure is schematically illustrated above with helices represented as rectangles and unstructed loop regions as lines.

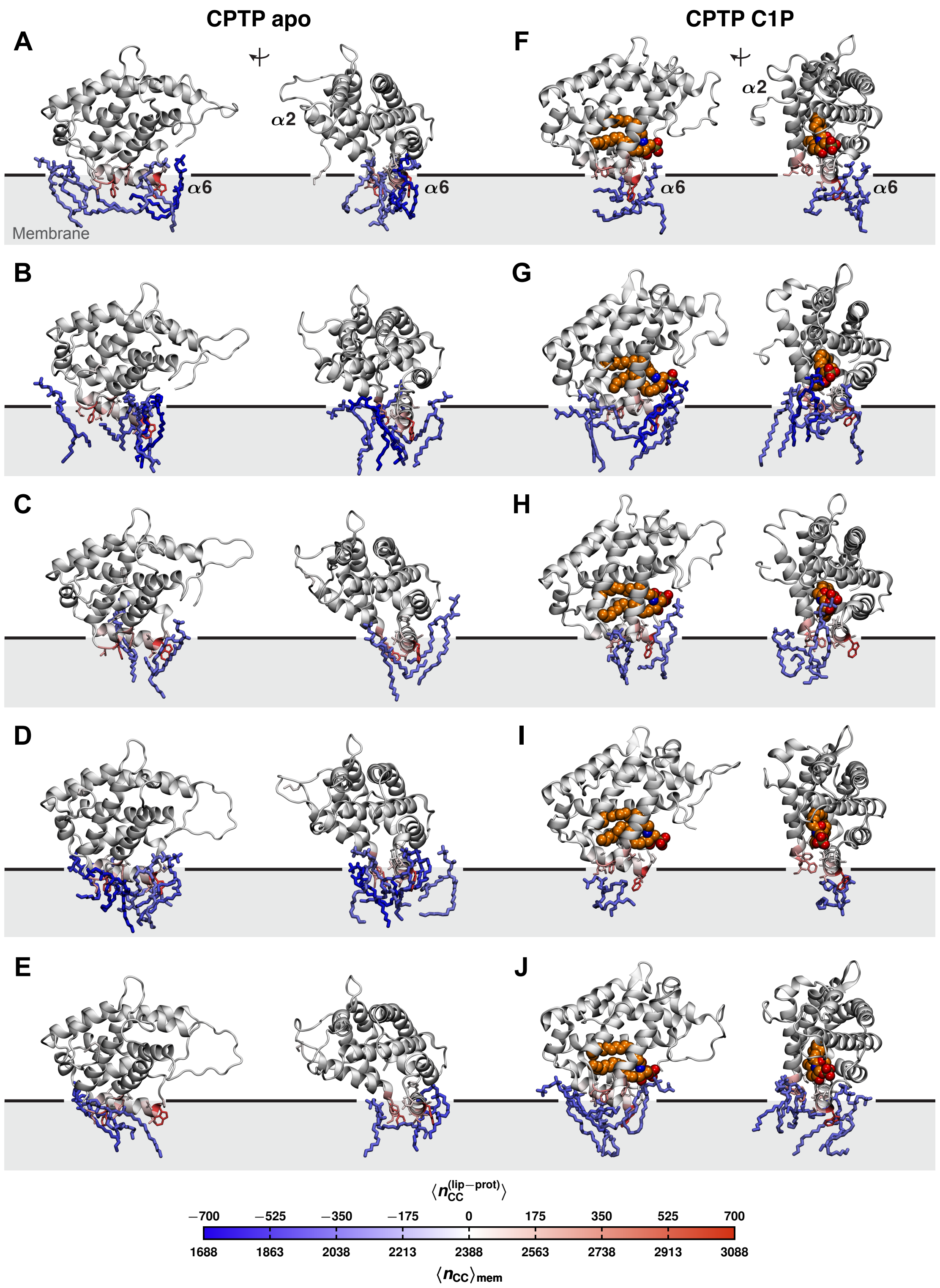


**Fig S7. CPTP forms hydrophobic contacts with lipids to disrupt their hydrophobic membrane environments.** $\langle n_{CC}^{(lip-prot)} \rangle$ mapped onto the structures of the (A-E) apo and (F-J) C1P-bound forms. Example configurations of lipids with (1) a reduction in $n_{CC}$ of $\geq$ 300 contacts relative to $\langle n_{CC} \rangle_{mem}$ of a POPC membrane without CPTP bound and (2) $\geq$ 90% of those contacts replaced with hydrophobic contacts with CPTP are shown. The black line indicates the average position of phosphate groups of membrane lipids.

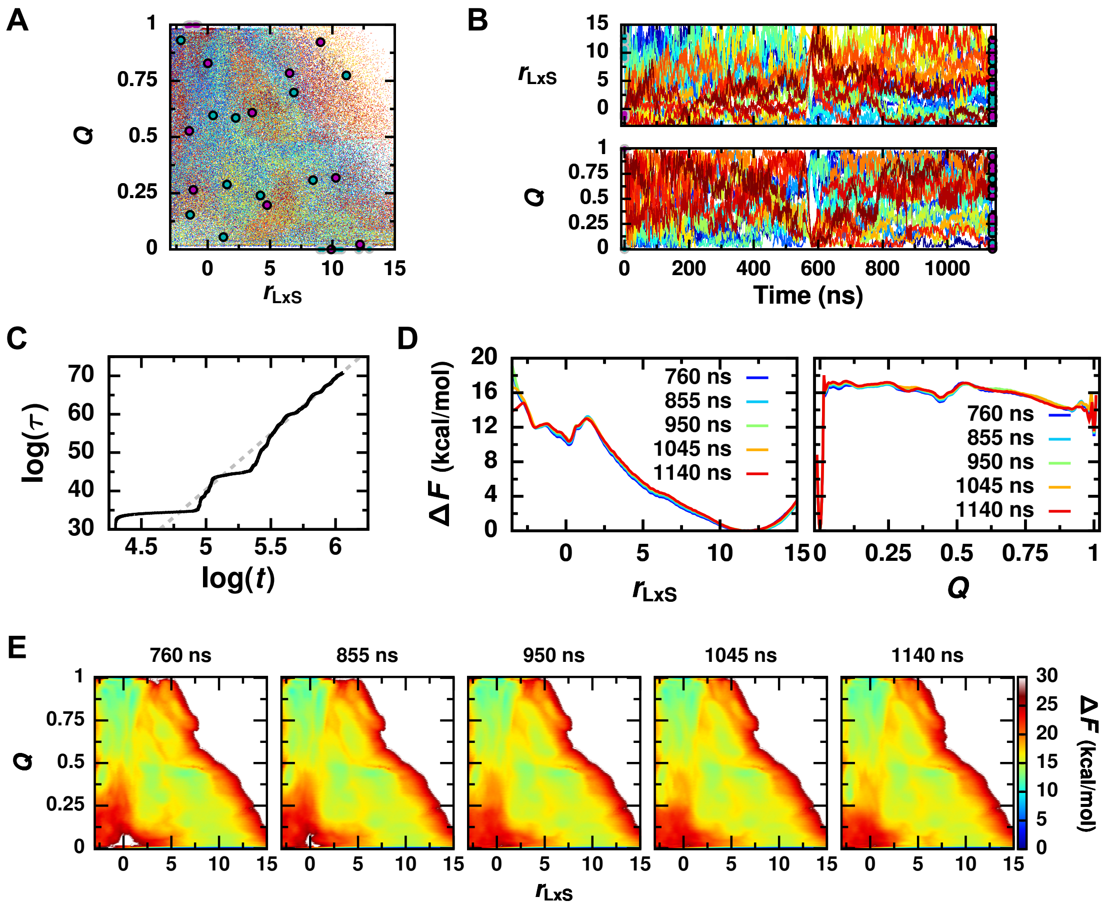


**Fig S8. Multi-walker well-tempered metadynamics simulation converges after 500 ns.** (A and B) Walkers initiated from the apo (cyan points outlined in gray) and C1P-bound forms (magenta points outlined in gray) broadly and diffusively sample the space of $r_{\mathrm{LxS}}$ and $Q$ and are well-mixed by the end of the simulation. Values of $r_{\mathrm{LxS}}$ and $Q$ at the end of the simulation are indicated with points outlined in black for each walker (cyan for those initialized from the apo form and magenta for those initialized from the C1P-bound form). (A) Values of $r_{\mathrm{LxS}}$ and $Q$ sampled during the simulation. Each point is colored by time with configurations sampled at early times shown in dark blue and at later times in dark red. (B) Trajectories of $r_{\mathrm{LxS}}$ and $Q$ for each walker. Walkers initiated from the apo form are plotted in cool colors, and walkers initiated from the C1P-bound form are plotted in warm colors. (C) Scaled time computed from the time-dependent bias is plotted versus simulation time on a log-log scale. The dashed gray line is the linear fit after 500 ns. The slope of this line is 29, which is consistent with the bias factor of 30 used in the simulation and suggests that a quasi-steady state has been reached. (D and E) Agreement between free energy profiles calculated at simulation times ranging from 760 ns (cumulatively 15.2 $\mu$s) to 1140 ns (cumulatively 22.8 $\mu$s) indicates convergence. All free energy profiles were calculated by reweighting frames after 500 ns (cumulatively 10 $\mu$s) up to the specified time using the estimator of Tiwary and Parrinello [91].

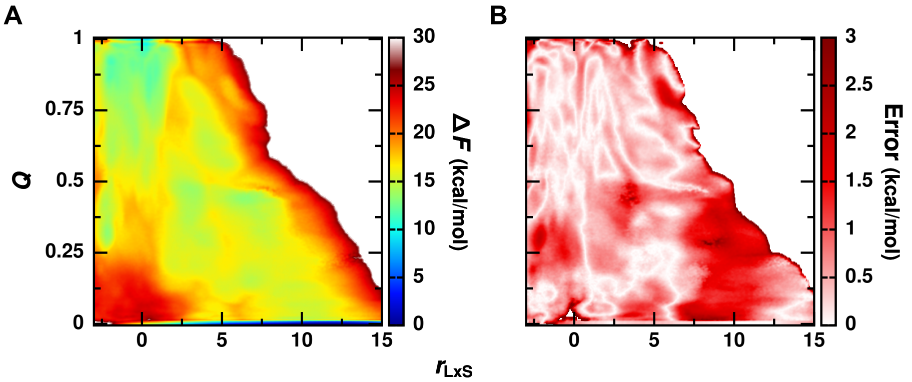


**Fig S9. Error in the free energy surface $\Delta F(r_{\mathrm{LxS}}, Q)$.** (A) The free energy surface $\Delta F(r_{\mathrm{LxS}}, Q)$ shown in Fig 7A is reproduced. (B) Standard error of $\Delta F(r_{\mathrm{LxS}}, Q)$ computed with block averaging.

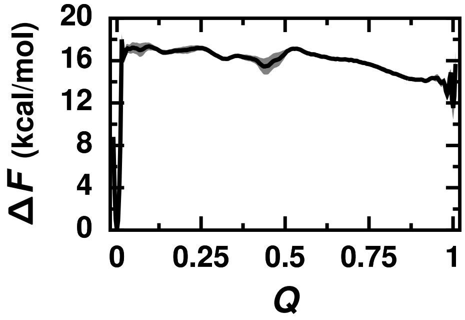


**Fig S10. Free energy profile as a function of $Q$.** $\Delta F(Q)$ is obtained by marginalizing $\Delta F(r_{\mathrm{LxS}}, Q)$ over $r_{\mathrm{LxS}}$. Error bars indicate the standard error computed with block averaging.

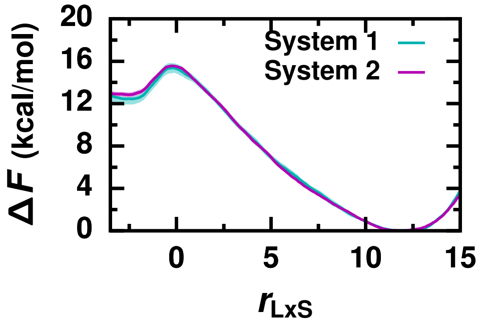


**Fig S11. Agreement between free energy profiles of passive C1P transport calculated for two independent systems indicates convergence of umbrella sampling simulations.** Free energy profiles along $r_{\mathrm{LxS}}$ calculated for two independent systems. For each system's free energy profile, error bars were calculated as the standard error of $\Delta F$ estimated from four independent 8 ns blocks.

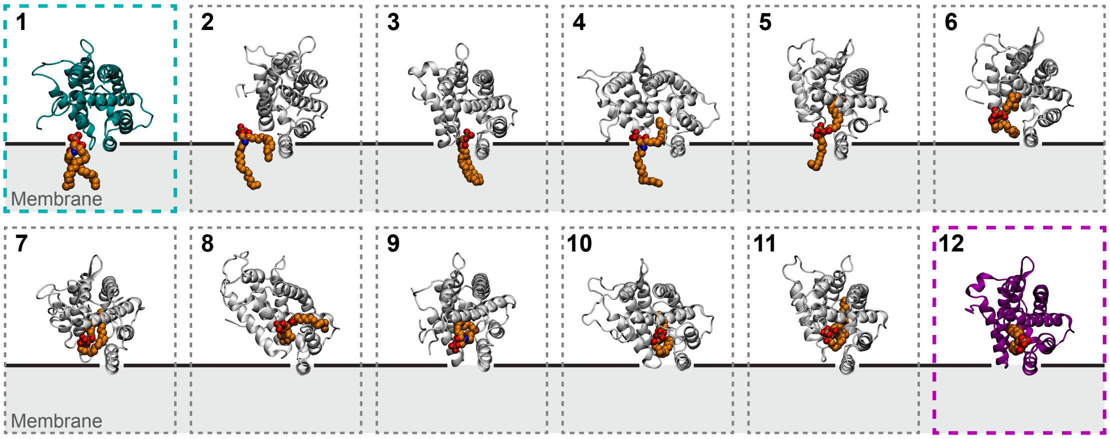


**Fig S12. Many different intermediate configurations are sampled during CPTP extraction (or insertion) of C1P from (or into) a membrane.** View looking into CPTP's hydrophobic cavity of example configurations from different regions of the free energy landscape outlined in Fig 7A. C1P is rendered as van der Waals spheres and colored orange. The apo form of CPTP in configuration 1 is colored dark cyan, and the C1P-bound form in configuration 12 is colored dark magenta. The black line indicates the average position of phosphate groups of membrane lipids.

### SUPPORTING NOTES

**Supporting Note 1: Principal component analysis of CPTP configurations.** We additionally characterize differences in the all-atom configurations of the apo and C1P-bound forms of CPTP both in solution and bound to the membrane through principal component analysis (PCA). Specifically, PCA was performed on frames from all simulations after alignment about C$\alpha$ atoms of helix $\alpha 6$, which approximates the plane of membrane surface, to characterize variations in the Cartesian coordinates of helices' backbone atoms. PCA was performed using GROMACS tools.

The first principal component (PC1) captures 48% of the total variation in the structures, and PCs $1-4$ collectively capture 89% of the total variance (Fig N1A). Fig N1B-D illustrate the structural differences described by PCs $1-4$ and the distributions of PC values sampled in solution-phase and membrane-bound simulations of both apo and C1P-bound forms of CPTP:

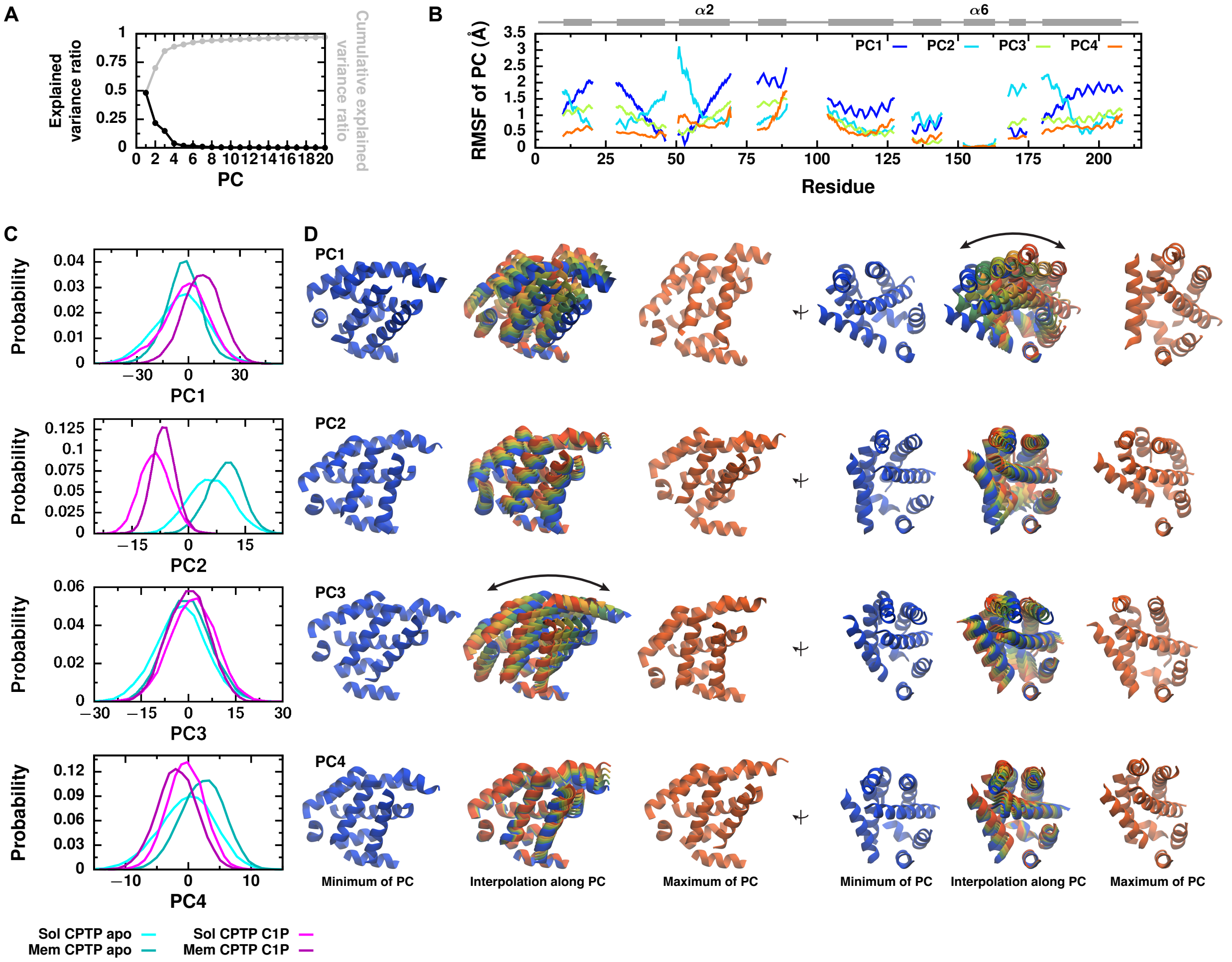


**Fig N1. Principal component analysis (PCA) captures differences in the all-atom structures of the apo and C1P-bound form of CPTP both in solution and bound to the membrane.** PCA was performed on helices' backbone atom's Cartesian coordinates after alignment about the C$\alpha$ atoms of helix $\alpha 6$. (A) Explained variance ratio of each PC and cumulative explained variance ratio. (B) Root-mean-square fluctuation (RMSF) of each atom along PCs $1-4$. CPTP's secondary structure is schematically illustrated above with helices represented as rectangles and unstructured loop regions as lines. (C) Configurations were projected on each PC, and distributions of the PC values are shown for the apo and C1P-bound forms of CPTP both in solution-phase and membrane-bound simulations. (D) Structural differences captured by each PC are illustrated by structural interpolations between the extreme projections along each PC.

PC1 describes a concerted rotation about helix $\alpha 6$, which approximates the surface of the membrane. Negative values of PC1 correspond to structures in which helix $\alpha 2$ is located closer and more parallel to the membrane surface, whereas positive values correspond to structures with helix $\alpha 2$ orientated more perpendicular to the membrane surface. We find that structures of the C1P-bound form of CPTP bound to the membrane have increased values of PC1 on average compared to either the C1P-bound form in solution or the apo form. Thus, membrane binding of the C1P-bound form promotes a concerted reorientation that aids the opening of gating helix $\alpha 2$ and that positions a widened entrance to CPTP's hydrophobic cavity at the membrane surface.

PC2 describes an internal reorganization of CPTP's helices that rotates the sides of its sandwich-like structure relative to each other (as if two stacked planar sheets were rotated relative to each other). Such changes captured by PC2 are evocative of a cleft-like gating mechanism (Fig 1 and S1 Fig) [16]. Of the first four PCs, variation along PC2 best captures differences between the apo and C1P-bound forms of CPTP, regardless of if CPTP is in solution or bound to the membrane. Indeed, C1P uptake (or release) results in substantial rearrangement of the sides of CPTP's sandwhich-like structure relative to each other (Fig 1 and S1 Fig). Membrane-bound structures of both the apo and C1P-bound forms of CPTP have increased values of PC2 on average compared to their respective solution-phase structures. Thus, membrane binding promotes a consistent change in the cleft to CPTP's hydrophobic cavity.

PC3 describes a concerted rotation orthogonal to that of PC1; if motion along PC1 were described as 'rocking side-to-side', then motion along PC3 would be described as 'rocking fowrard-and-backward'. Conformational ensembles of the apo and C1P-bound forms sampled in both solution-phase and membrane-bound simulations exhibit similar variation along PC3.

PC4 describes an internal reorganization of CPTP's helices different from that of PC2. While solution-phase structures of CPTP have similar average values of PC4, the membrane-bound structures of the apo and C1P-bound forms have average values different from each other and from the solution-phase structures.

Overall, PCA indicates that structures of the apo and C1P-bound forms differ both in solution and bound to the membrane, and that membrane binding can promote opening of gating helix $\alpha 2$ and conformational changes suggestive of a cleft-like gating mechanism.

**Supporting Note 2: Definition of $Q$.** The fraction of contacts C1P makes with CPTP when fully inside its hydrophobic cavity, $Q$, was used as a second order parameter (or collective variable) for biased simulations. $Q$ was chosen to reliably identify configurations with C1P inside CPTP versus configurations with C1P outside CPTP and to enhance the sampling of CPTP–C1P interactions. Since $r_{\mathrm{LxS}}$ only describes C1P–membrane interactions, it accomplishes neither of these things, while $Q$ does. Specifically, CPTP–C1P contact pairs used to calculate $Q$ were selected to capture:

1. Hydrophobic contacts between carbons of C1P and carbons of residues lining CPTP's hydrophobic cavity. Carbon-carbon (CC) pairs were selected based on their average distance, $d_{\mathrm{C1P-CPTP}}$, in solution-phase simulations of the C1P-bound form of CPTP (Fig N2A). To minimize the computational expensive of calculating $Q$ during biased simulations, which increases with the number of CC pairs considered, while accurately identifying configurations with C1P inside CPTP's hydrophobic cavity, a cutoff of $d_{\mathrm{C1P-CPTP}} \leq 7.8$ Å was used to select these CC pairs. Residues with these CC pairs used to calculate $Q$ are shown in Fig N2C.

2. Polar contacts between C1P's headgroup and sphingoid backbone and residues at the entrance to CPTP's hydrophobic cavity. Heavy atom pairs were selected based on $d_{\mathrm{C1P-CPTP}}$ (Fig N2B). To minimize the computational expense of calculating $Q$ during biased simulations while capturing all important polar contacts, a cutoff of $d_{\mathrm{C1P-CPTP}} \leq 5.5$ Å was used to select these pairs. Residues with these polar atom pairs used to calculate $Q$ are shown in Fig N2D.

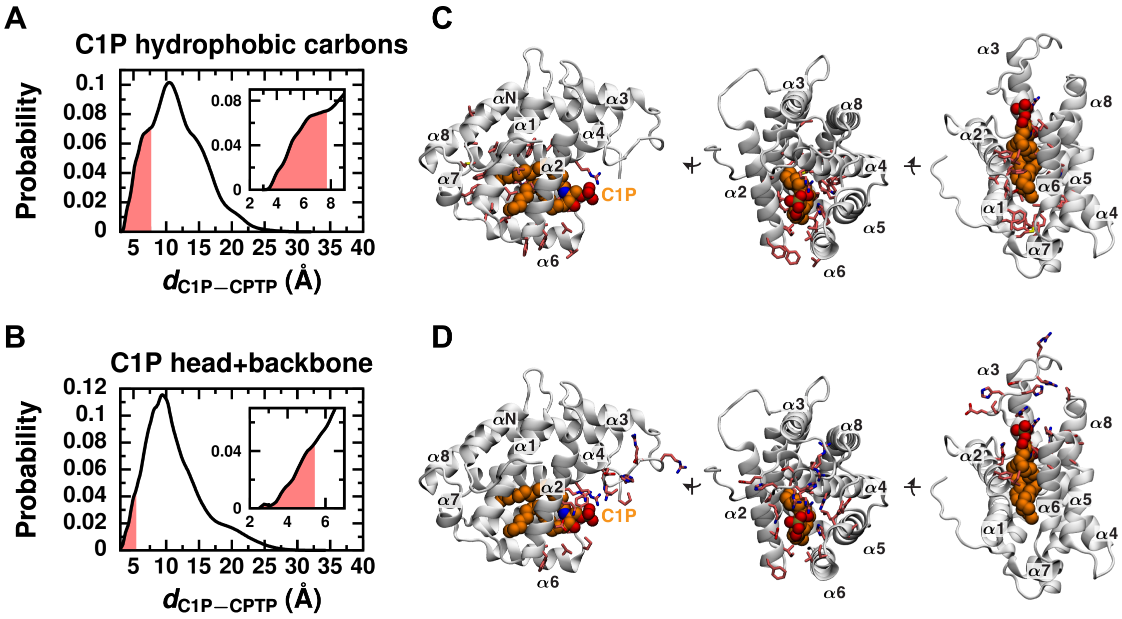


**Fig N2. CPTP–C1P contacts used to define $Q$ account for key hydrophobic and polar interactions.** (A and B) Distribution of the average distance, $d_{\mathrm{C1P-CPTP}}$, between (A) carbon-carbon pairs and (B) pairs of polar atoms of C1P and CPTP during all-atom solution-phase simulations of the C1P-bound form of CPTP. Carbon-carbon pairs with $d_{\mathrm{C1P-CPTP}} \leq 7.8$ Å are used to calculate Q and highlighted by the red region in (A). Residues with these carbon-carbon pairs are rendered in licorice and colored red in (C). Pairs of polar atoms with $d_{\mathrm{C1P-CPTP}} \leq 5.5$ Å are used to calculate Q and highlighted by the red region in (B). Residues with these pairs of polar atoms are rendered in licorice and colored red in (D).

Based on these criteria for selecting contact pairs, 1,176 atom pairs are used to calculate $Q$ using Eq. 4 given in the Methods.

We confirmed that $Q$ reliably identifies configurations with C1P fully inside CPTP's hydrophobic cavity and orientated as in crystal structures from configurations with C1P outside, partially inside, or improperly orientated within CPTP's cavity. To do so, we monitored the value of $Q$ during all-atom simulations in which C1P enters into CPTP's hydrophobic cavity from the solvent. While the relaxation process that occurs during these simulations is not cellularly relevant, it is sufficiently rapid to observe in unbiased simulations. Thus, we are able to harvest multiple all-atom trajectories of C1P entry from solvent and use them to benchmark $Q$. Five simulations, each initialized with C1P randomly placed in the solvent around the apo form of CPTP, were performed using the same parameters as the all-atom solution-phase simulations described in the Methods. Each simulation was run for a maximum of 2 $\mu$s or until both tails of C1P were inserted into CPTP and no longer exposed to solvent based on visual inspection.

Fig N3 shows the value of $Q$ during each of these simulations and the final configuration of C1P bound to CPTP. In three of the simulations (shown in green, cyan, and blue in Fig N3), C1P inserts between the seam

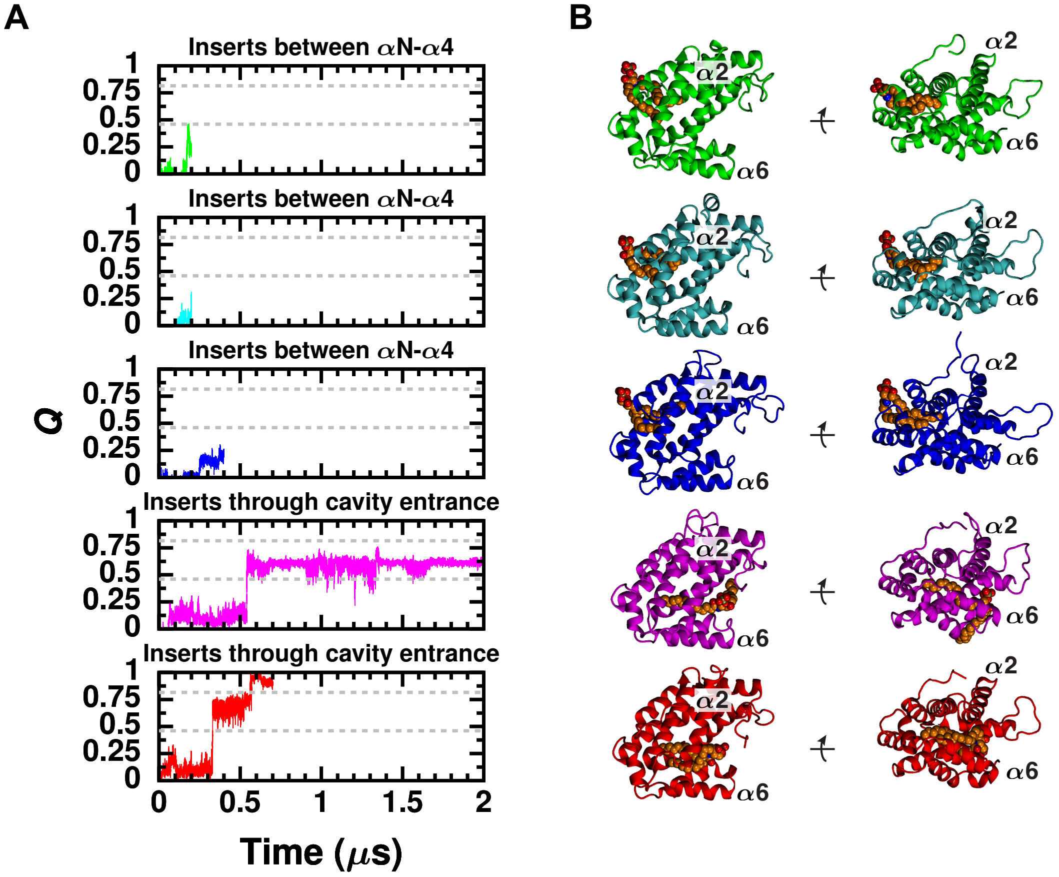


**Fig N3. Analysis of unbiased simulations of C1P entry into CPTP's hydrophobic cavity from solvent indicate that $Q$ reliably distinguishes configurations with C1P fully inside CPTP's hydrophobic cavity from others.** (A) Value of $Q$ during five independent simulations of C1P entry from solvent. In the green, cyan, and blue trajectories, C1P enters through the top seam between helices $\alpha$N and $\alpha$4. In the magenta and red trajectories, C1P enters through the entrance identified in crystal structures and located at the membrane surface in simulations. The dashed lines mark the values of $Q$ where sharp transitions occur in the red trajectory. (B) Final configurations of each simulated trajecotry.

created by helices $\alpha$N and $\alpha$4. These trajectories are not representative of C1P uptake from a membrane since insertion through helices $\alpha$N and $\alpha$4, which are fully exposed to solvent when CPTP is bound to a membrane (Fig 2), would require C1P to become fully solvated before entering CPTP's hydrophobic cavity. Thus, they serve as valuable tests of using $Q$ to accurately identify configurations with C1P properly housed in CPTP's cavity. $Q = 1$ when C1P is properly housed inside CPTP's cavity, whereas $Q$ never surpasses 0.5 in these trajectories. In the other two trajectories (shown in magenta and red in Fig N3), C1P inserts through the entrance to CPTP's hydrophobic cavity as occurs when it's extracted from a membrane. In both trajectories, C1P's tails enter individually. In the trajectory shown in magenta in Fig N3, the second tail fails to enter within 2 $\mu$s. In the trajectory shown in red in Fig N3, both tails enter within 700 ns. In both trajectories, $Q$ rapidly changes from $Q \approx 0.1$ to $Q \approx 0.6$ when the first tail enters, and, in the red trajectory, then changes rapidly again when the second tail enters. Thus, $Q$ distinguishes different ways that C1P can bind to CPTP and can be used to enhance the sampling of interactions between C1P and CPTP. We note that these trajectories do not provide any evidence that $Q$ is the reaction coordinate [32–34] (or necessarily a component of the reaction coordinate) for CPTP-mediated C1P transport.
